# Differential Expression Analysis for Spatially Correlated Data

**DOI:** 10.1101/2024.08.02.606405

**Authors:** Ana Gabriela Vasconcelos, Daniel McGuire, Noah Simon, Patrick Danaher, Ali Shojaie

## Abstract

Differential expression is a key application of imaging spatial transcriptomics, moving analysis beyond cell type localization to examining cell state responses to microenvironments. However, spatial data poses new challenges to differential expression: segmentation errors cause bias in fold-change estimates, and correlation among neighboring cells leads standard models to inflate statistical significance. We find that ignoring these issues can result in considerable false discoveries that greatly outnumber true findings. We present a suite of solutions to these fundamental challenges, and implement them in the R package smiDE. spatial transcriptomics, differential expression, segmentation error mitigation, spatial correlation, spatial random effects model

## 1 Introduction

Imaging-based spatial transcriptomics (ST) technologies detect tissues’ RNA molecules in situ, producing single cell expression data while preserving cell’s spatial locations [1, 2]. This pairing of gene expression data with spatial context creates an opportunity to survey how cells respond to their environment or to interactions with other cell types. Consider, for example, a tumor containing cancer cells, T-cells, and other cell types (Figure 1a) falling in two distinct regions: the tumor bed and the stroma (Figure 1b). Spatial transcriptomics allows us to answer questions such as: how do one or more cell types modulate their gene expression differently across functional regions of the tissue (Fig 1c)? Or, how does it change gene expression when co-localized with another cell type (Fig 1d)? Or how does the cell type change expression in treated vs. untreated tumors?

**Figure 1:**
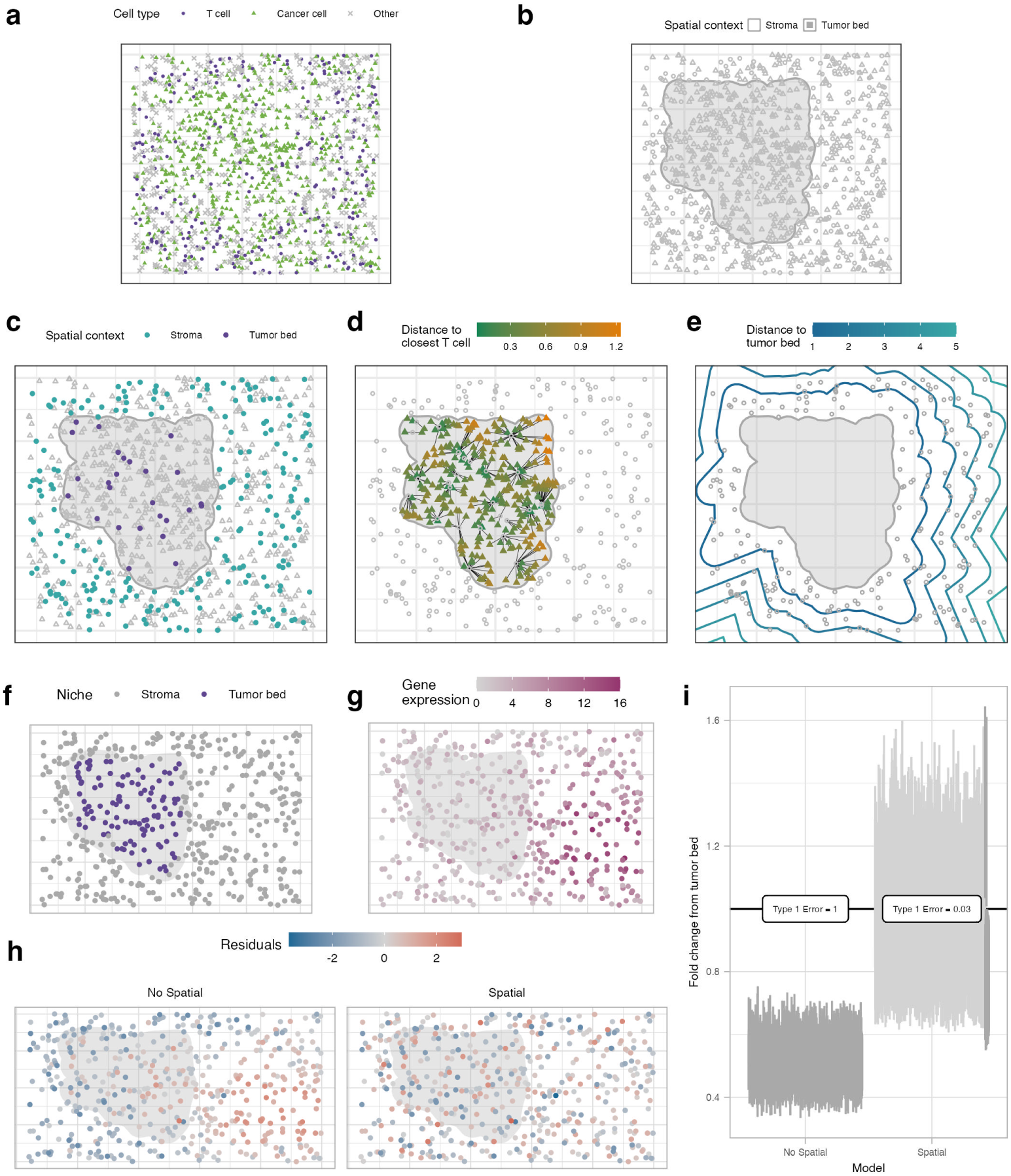
Examples of possible DE tests in a spatial context (a-e) and toy simulation to illustrate spatial confounding(f-i). Assume we are interested in T-cells and cancer cells **(a)**, which are located either in a tumor bed or stroma **(b)**. Possible examples of DE analysis are: contrast gene expression between T-cells in tumor bed vs T-cells in stroma **(c)**; analyze how expression of genes of cancer cells in tumor bed differentiates based on distance to nearest T-cell **(d)**; or determine how expression of genes of T-cells in stroma relates to distance to tumor bed **(e)**. For the simulation, **f**, cells are located in two different spatial context, tumor bed and stroma. **g**, shows gene expression level in each cell. Data was generated unrelated to the region, but with a spatial pattern to it. **h**, residuals obtained under a model that assumes data is independent and thus does not adjust for spatial, and another that accounts for spatial correlation using a Gaussian Process. We observe that there is a pattern in the residuals of the independent case, which indicates the lack of fit of the model, whereas in the spatial model the pattern is not as present. **i**, shows the confidence interval for the fold change estimate of stroma compared to tumor bed in each model. We see that for the independent case none of the CIs cover the true value of one due to bias, that is, under that model there is an indication that the gene is differentially expressed between niches, whereas the model considering the spatial correlation has better control of type 1 error.

These questions can be answered through differential expression (DE) analysis. DE analysis has been a mainstay of gene expression studies, and it becomes even more powerful in the context of single cell spatial transcriptomics. Many methods for DE analysis are available, such as DEseq2 [3], limma [4], and MAST [5], which were originally developed for traditional bulk or single cell gene expression studies. However, spatial transcriptomics data presents unique challenges that these methods were not designed for. First, imperfect segmentation of cell boundaries means some RNA molecules are incorrectly assigned to neighboring cells, inducing spatially-dependent bias in expression profiles. Second, cells that neighbor each other are likely to have correlated expression profiles. This correlation can be thought of as cells’ response to spatial variables omitted from the differential expression model; these variables act as potentially powerful confounders which, if ignored, can badly distort differential expression results (Figure 1f-i). Finally, sample sizes can be enormous, with *>* 10^6^ cells assayed per sample. While this problem is not unique to spatial transcriptomics, it makes it computationally expensive to model the spatial correlation between cells.

Despite their importance, the main challenges of DE analysis in spatial transcriptomics have not been fully addressed. For instance, C-SIDE, which is a common method used for DE [6], focuses on handling spot-based data with multiple cells and adjusts for proportions of cell types. While C-SIDE addresses the challenge of cell segmentation uncertainty, as with other existing approaches for DE in this context, it fails to accurately account for the spatial variation of genes. More recently, Ospina et al. [7] have also discussed the importance of accounting for spatial correlation to achieve better model fit and avoid false positives. However, their use of existing methods for Gaussian observations may not account for excess zeroes. Moreover, their methods remains computationally intensive and are not benchmarked; therefore it is unclear whether they control the type 1 error and provide sufficient power. On the other hand, methods that account for spatial variation, such as SpatialDE [8], SPARK [9], Trendscreek [10] and BOOST-GP [11], focus on identifying genes with *spatially variable expression* (SE). These methods perform variance component tests that decompose the variability between spatial and non-spatial sources. However, instead of finding genes with expressions that vary across space, many studies require finding genes differently expressed (DE) between regions [12, 13], or cell types [14], which corresponds to a different goal; this problem is the focus of the current study.

In this paper, we propose solutions to the major challenges of differential expression with spatial data. Through simulation and real data applications, we show the importance of accounting for these issues to avoid false discoveries and achieve more reproducible and generalizable findings. We present two countermeasures to deal with cell segmentation errors, and benchmark several solutions for dealing with spatial correlation, as well as develop a novel variant of spatial random effects models that addresses the challenges of spatial correlation while improving computational efficiency. A software with all the implementations is provided, as well as recommendations for best practices.

## 2 Results

### 2.1 Choosing parametric distribution for DE regression model

For scRNA-seq data, Negative Binomial regression models are commonly used to model the observed expression of RNA transcripts across cells, often using random effects in multi-subject studies to account for heterogeneity across independent biological samples [15].

Assuming the data are generated from a Negative Binomial distribution is a reasonable a priori assumption, as count data are often overdispersed relative to a Poisson distribution due to both technical and biological sources of variation [16]. Due to the increasing scale of modern ST datasets, with potentially millions of cells across multiple biological samples, we compared performance tradeoffs of the principled choice of a reference Negative Binomial regression model against approximations which may be more computationally tractable in large-scale studies.

A Negative Binomial mixed model with random effects (RE) predicted for biological samples (Negative Binomial RE) may be chosen from first principles as the model that is expected to best describe the data. We may specify the regression model as

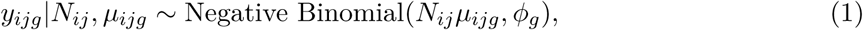

where *y_ijg_* are the observed counts for cell *i*, biological sample *j*, and gene *g*. *N_ij_* are the total observed transcripts for cell *i* across all genes, and *µ_ijg_* is the expected rate of expressed transcripts for cell *i*, gene *g*. Parameter *ϕ_g_* is a size factor which tracks with the level of overdispersion in the expression of gene *g* relative to a Poisson distribution. The expected expression rate *µ_ijg_* may be parameterized by covariates

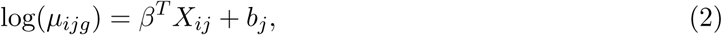

where *X_ij_* and *β* represent the observed covariates of interest associated with cell *i* in sample *j* and their regression coefficients. The term *b_j_* is a random effect shared by all cells belonging to sample *j*.

As an alternative to the Negative Binomial RE model, we also consider a Poisson model (Poisson RE) parametrized as

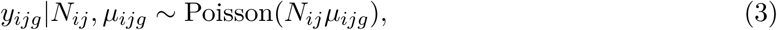

with the same parameterization for *µ_ijg_* as in (2). Finally, we also consider a simple Gaussian model (Gaussian RE). Due to the importance of the total transcript count *N_ij_* as a cell specific scaling term which accounts for technical variation in cells’ measurement efficiency, and the desire to compare fold change estimates on the same scale as the log-linear Negative Binomial and Poisson models, we consider the following normalization for regression with Gaussian family:

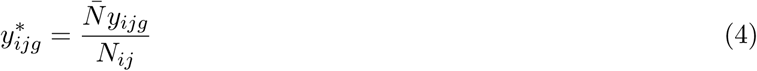

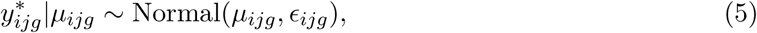

where 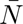 is the average total transcript count across all cells. For the Gaussian RE case, we avoid a direct log transformation of the normalized counts. Use of a log transformation would require taking the form log(*y_ijg_* ∗ +*c*), where a constant ‘*c*’ must be chosen to avoid infinite values when the raw count is zero. In this transformation, the choice of ‘*c*’ is arbitrary and can potentially have a non-negligible impact on the regression model and its error distribution, particularly for low expressed genes where zero counts are frequent.

Using the CosMx NSCLC dataset [1] with 487,819 cells from 5 NSCLC tissues, we compared the above parametric models (Negative Binomial RE, Poisson RE, and Gaussian RE) in terms of computation time, concordance in effect size estimation, and significant gene calls. We ran DE regression models for 7 cell types (macrophage, T-reg, T-CD8, T-CD4, plasmablast, neutrophil, and tumor). A categorical covariate ‘niche’ annotates the functional region of tissue the cell resides in, and was treated as the primary covariate of interest in the regression models. This ‘niche’ covariate had 9 categories (tumor interior, storma, tumor-stroma boundary, myeloid-enriched stroma lymphoid structure, immune, plasmablast-enriched stroma, and macrophages). For each cell type, we assessed whether a gene was DE in a particular niche compared to the cell-weighted average of all other niches.

We found that concordance in fold change estimates between all three parametric families was high, with fold change estimates between Negative Binomial and Poisson families virtually identical in macrophage cells (Fig 2a), and across cell types (Supp Fig 1). Poisson RE models produced consistently larger test-statistics (in magnitude) and smaller p-values compared to Negative Binomial and Gaussian RE models (Fig 2b). Such inflated type I errors are expected from models that fail to account for count overdispersion, resulting in standard errors which are too small. Gaussian regression tended to call fewest significant genes of any family in macrophages (Fig 2b,c) and across cell types (Supp Fig 2); however, a higher proportion of Gaussian RE significant gene-contrasts overlapped with Negative Binomial RE (238/284 = 83%) compared to Poisson RE (320 / 408 = 78%) (Fig 2c), a pattern which persisted across cell types (Supp Fig 3). We observed that the degree of overlap in significant gene sets between families tended to increase with the number of cells analyzed (Supp Fig 3).

**Figure 2:**
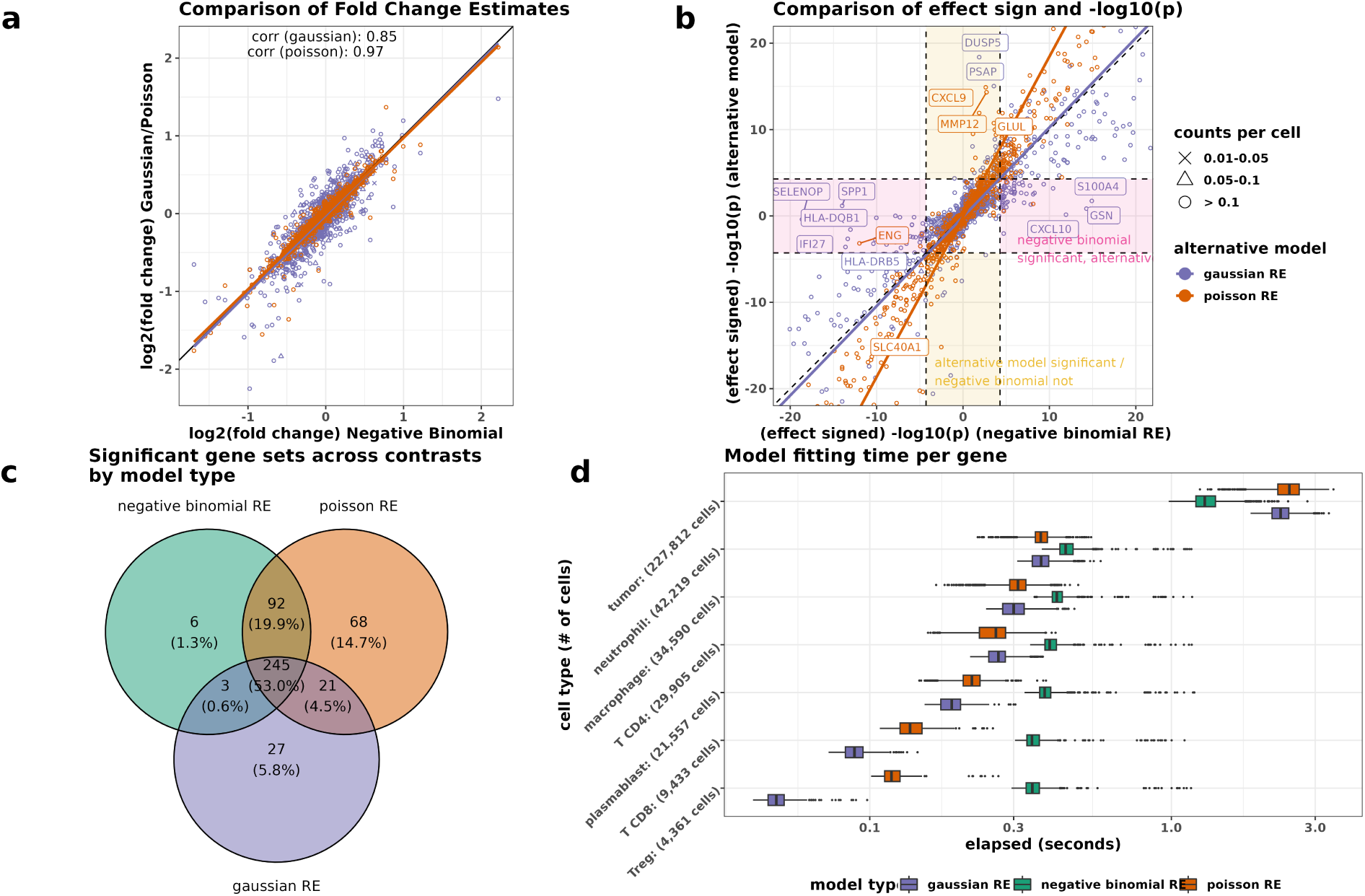
Comparison between parametric families for cell type specific DE between spatial niches. **a**, log2 fold changes for genes DE in macrophage cells between negative binomial RE models (x-axis) compared to the corresponding log2 fold change estimates in gaussian or poisson RE models. **b**, -log10(p-values) multiplied by the estimated direction of effect (+/-) compared between negative binomial RE models (x-axis) with gaussian or poisson RE models (y-axis) for macrophage cells. Highlighted pink regions indicate genes where a spatial contrast and gene is significant using negative binomial RE, and but not significant using alternative model at a bonferroni threshold of (0.05/(total # of genes in the panel)). **c**, Venn diagram of overlapping significant gene contrasts between 3 model types. **d**, Computation time comparisons between gaussian, negative binomial, and poisson RE models across 7 cell types.

In terms of computation efficiency, Gaussian RE models were fastest in elapsed time per single gene when the number of cells modeled were small, though the gap in modeling time became smaller as the number of cells increased. Model estimation time was similar overall at roughly 30-40k cells (Fig 2d). Somewhat surprisingly, Negative Binomial RE models were fastest at the largest sample size of DE for 227k tumor cells. Negative Binomial and Poisson RE models were estimated using the NEBULA package, which implements a fast large sample analytical approximation to the model estimation, resulting in computation times which are orders of magnitude faster than other package implementations [15]. However, NEBULA does not support analysis of spatially correlated random effects, so the optimal models discussed in *Spatial Modeling* do not benefit from the speed described here.

### 2.2 Countermeasures for cell segmentation error

Often a key challenge in cell type specific analysis of ST data is dealing with the impact of erroneous transcript assignment due to imperfect or overlapping cell boundaries. While many methods have been developed for single cell segmentation of ST data [17, 18], even minor errors may potentially confound or distort cell type specific downstream analyses. As an illustrative example, we consider an analysis of macrophage cells which seeks to identify genes DE in tumor cell infiltrated regions (tumor-interior) of the tissue compared to the rest of the tissue. Identifying genes in macrophage cells that are DE in the tumor-interior may reveal important biological mechanisms by which macrophages promote immunosuppression or regulate cancer progression in the tumor [19, 20, 21]. Figure 3a shows a macrophage cell surrounded by cancer cells in a tumor bed. Due to imperfect definition of cell boundaries, a small number of KRT17 transcripts expressed by a neighboring cancer cell are falsely assigned to the macrophage cell. In the DE model, these errors are correlated with our scientific question in a way that confounds the DE analysis (Fig 3b); the tumor-interior region is more likely to have KRT17-impacted macrophage cells than non-tumor regions of tissue due to the abundance of surrounding cancer cells—the abundance of cancer cells being a primary signature of the ‘tumor-interior’.

**Figure 3:**
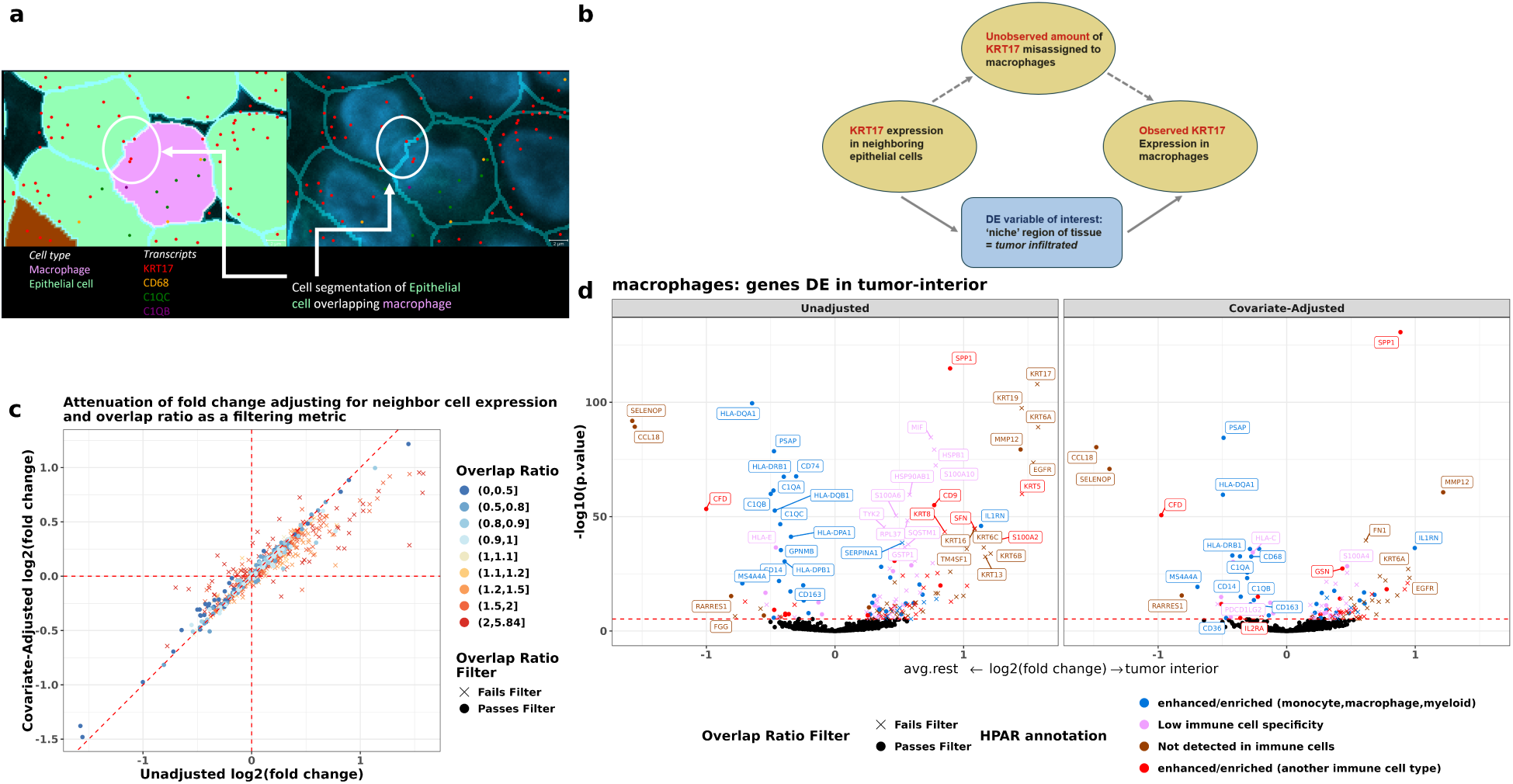
Impact of segmentation error on DE analysis. **a**, Illustrative example of a macrophage cell (pink) surrounded by cancerous cells in a tumor cell-infiltrated region of tissue. KRT17 transcripts (red dots) expressed by a neighboring epithelial cell breaches the estimated border of the macrophage cell. The cell borders are ambiguous; evident by overlapping nuclei of the two cells when viewed in two dimensions result in false positive counts of KRT17 assigned to the macrophage. **b**, DAG showing causal path by which segmentation error correlates with both the response (KRT17 expression in macrophage) and estimand of interest (DE variable), acting as an unobserved confounder. Covariate adjustment via KRT17 expression in neighboring tumor cells can act as a proxy variable to reduce confounding bias. **c**, Estimates of log2 fold change, with and without covariate adjustment for neighboring cell expression in macrophages. Genes which are more expressed in neighboring cell types (red, Overlap Ratio *>* 1), are attenuated towards zero after adjustment, while genes which are expressed more in the cell type of interest (blue, Overlap Ratio *<* 1) have similar estimates with and without covariate adjustment. **d**, A comparison between ‘Unadjusted’ and ‘covariate-adjusted’ volcano plots. Each gene is colored by a condensed description of the annotation provided for “Immune cell specificity” in the Human Protein Atlas (HPAR)[22, 23]. Based on prior biology, blue annotated genes are known to be expressed in the analyzed cell type (macrophage) and are the least controversial while brown genes (“Not detected in immune cells”) are the most prone to be suspect. The majority of significant suspect genes become non-significant in the adjusted model. Some remaining suspect brown genes may be further disregarded in an analysis which applies a gene filtering metric.

Next, we introduce two countermeasures to the artifacts caused by segmentation errors. First, we propose a rule for flagging genes within a celltype most likely to be impacted by segmentation errors. Second, we expand our models to explicitly model gene expression in neighboring cells as a control variable, thereby weakening the effect of segmentation error as a confounder.

#### A segmentation overlap ratio metric for cell type-specific gene filtering

We motivate a metric for preemptively excluding genes likely to return spurious DE results due to imperfect or overlapping cell segmentation boundaries, which we call an “overlap ratio metric”. Intuitively, when analyzing a specific cell type, we expect that a gene may be prone to error if it has low expression in the cell type of interest and high expression in cells spatially adjacent to the cell type of interest (Fig 3a).

Let 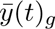 denote the average expression of a particular gene *g* for the type *t* cells of interest. Further denote the average expression per cell of the gene in the *neighboring cells* of other cell types by 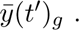 Then the we define the overlap ratio metric as

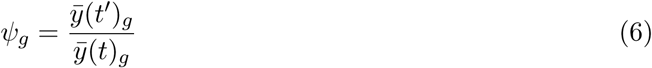

This metric quantifies susceptibility to confounding bias from overlapping or erroneous cell boundaries. Genes with *ψ_g_ >* 1 are higher expressed, on average, in neighboring cells of other types than that of the cell type of interest; hence the observed number of transcripts in analyzed cells may contain unacceptable levels of error. While values other than 1 might be considered as cutoffs for determining whether to analyze a particular gene for a given cell type, we use the threshold of 1 as an intuitive cutoff in our analysis (Fig 3 c,d). The optimal cutoff will vary with the accuracy of cell segmentation and with the analyst’s tolerance for false discoveries. Regardless of the cutoff, this statistic provides a measure of the skepticism each gene deserves. A detailed formulation is provided in Methods.

#### A covariate adjustment approach using gene expression in neighboring cells

An alternative or complementary approach to removing a gene from analysis via overlap ratio metric, is to adjust for propensity of transcript assignment error within the regression model. In practice, the exact amount of erroneous transcripts in individual cells is unknown, but is correlated with the expression of the analyzed gene in spatially neighboring cells, making ‘total neighbor expression’ a useful proxy covariate to include in the regression model (Fig 3b).

Consider a gene and a set of cells of the same type. To capture the gene’s propensity for error within the analyzed cells, we measure its total expression in neighboring cells of other cell types (we expect that error from neighbors of the same cell type will have negligible impact on analyses where neighboring cells have highly similar values of predictor variables). Denote this propensity for error in transcript assignment with a spatially lagged expression vector *l_g_*, where *l_g_* = *W* (*r*)*y*(*t*^′^)*_g_*; here, *W* (*r*) is a neighborhood indicator matrix indicating the neighboring cells of other cell types within a distance *r* of the cells to be analyzed, and *y*(*t*^′^) is the expression of the gene in cell types *other than* type *t*. If we assume that errors in transcript assignment increase the average observed expression by a factor proportional to *l_g_*, we can update our regression model in Eq (2) to include this factor as a proxy covariate for unobserved transcript assignment error. The updated model is specified as

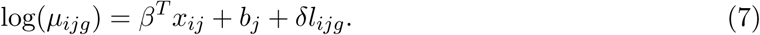

While we describe the expression in neighboring cells as having an additive effect, the inclusion of this covariate in log-linear regression models with a multiplicative effect can be a convenient and effective approximation.

Given that extremely close neighboring cells may be more likely to impact a cell of interest compared to farther neighboring cells, we may also choose to weight the neighboring expression by the distance of the neighboring cell, such that the elements of the neighborhood matrix *W* (*r*) follow

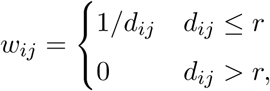

where *d_ij_* is the distance between a cell *i* to a neighbor cell *j*. To stabilize model estimation with this covariate (which is typically heavily right-skewed) we apply a rank-based inverse normal transformation to *l_g_*, i.e., 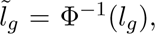 where Φ denotes the cumulative distribution function for a standard normal distribution. In practice, *l_g_* may be a noisy approximation of actual transcript assignment error, but a useful proxy variable, which may greatly reduce false positives while limiting the number of genes that must be disregarded via a gene filtering metric [24].

#### Covariate adjustment and gene filtering effectively mitigate false positive associations arising from segmentation error

We compared the behavior of the gene flagging metric and covariate adjustment method across DE analyses of macrophage, T-CD8, T-CD4, and T-reg cell types across 5 patients in the CosMx NSCLC dataset [1]. For each cell type, we fit Negative Binomial RE regression models and looked at genes DE in any of the 9 spatial niche regions annotated (Fig 3c,d). For each cell type, we considered both a naïve model without covariate adjustment for neighboring gene expression, as well as a covariate adjusted version, and considered the significant genes before and after applying a overlap ratio metric filter (Fig 3d).

To quantify the impact of segmentation error, we labeled genes according to their Human Protein Atlas Reference annotation for “Immune Cell Specificity” [22, 23]. Significant DE genes labeled as “Not detected in immune cells” are likely false positives that arise from segmentation errors, while genes labeled as “Enhanced or enriched” in the corresponding cell type may be considered plausible. Fig 3d compares volcano plots for macrophage cell DE genes in the tumor interior, where the naive unadjusted model identifies a number of cancer cell enriched keratin genes, which are over-expressed in the tumor-interior due to errors in segmentation. Fold changes and p-values for false positive genes were largely attenuated by covariate adjustment, and almost entirely eliminated when removing genes according to the overlap ratio filtering metric (Fig 3c,d). For example, likely false positive genes were removed for 112/138, 68/72, 27/27, and 14/16 significant genes labeled as “Not Detected in Immune cells” in macrophage, T-CD4, T-CD8, and T-reg celltypes, respectively, when using a combined approach of covariate adjustment and gene filtering (Fig 3d, Supp Fig 4).

### 2.3 Spatial modeling

Spatial transcriptomics techniques offer the advantage of preserving the cell organization within tissue samples. This attribute can be crucial since we expect that neighboring cells should have more similar gene expression profiles compared to cells located at greater distances. The importance of accounting for the correlation between cells is illustrated in the simulated example in Figure 1f-i.

The interest is to study if the expression of a gene differs in cells of the same type, say T-cells, that are located either on a tumor bed, or in the stroma (Figure 1f). Two modelling approaches are considered: one that accounts for spatial correlation (Gaussian Process explained further in Methods) and one that assumes independence of cells (No Spatial). In this example, gene expression in some cells is higher due to various spatially smooth factors unrelated to niche (Figure 1g), for example localized inflammation, hypoxia or cellular necrosis. These factors lead to correlation among gene expression in neighboring cells. Examining the residuals of the two models (Figure 1h) reveals a spatial pattern when not adjusting for the spatial correlation, which indicates lack of model fit. This lack of fit is mitigated when the spatial correlation is modeled. By not accounting for this correlation when testing for differential expression between T cells in tumor bed or in the stroma, we would incorrectly conclude that the gene is downregulated in T cells inside the tumor (Figure 1i), when in reality the difference in expression is related to other unrelated factors. This simple example highlights the importance of accounting for spatial correlation in order to avoid the spatial bias and correctly understand the true underlying signal.

#### Statistical approaches to dealing with spatial correlation

Not necessarily in a transcriptomics context, several classical spatial statistics models have been developed to deal with spatial correlation; these include Gaussian Processes [25], iCAR [26] or BYM2 [27]. Even though these models are well known and widely used in other spatial data applications, they have not yet been used for DE analysis with spatial single cell transcriptomics data. Therefore, we next introduce the Gaussian Process and BYM2 models in this context. Let *y_ig_* be the transcript counts of the *g*th gene (*g* = 1*, . . . , p*) on the *i*th cell (*i* = 1*, . . . , n*). Then for each gene *g* we model the conditional mean *µ_g_* of *y_g_* = (*y*_1_*_g_, . . . , y_ng_*) as

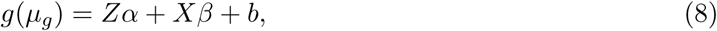

where *g* is a link function, either the identity for Gaussian distribution, or log function for Negative Binomial. The vector *X* represents the covariate of interest, with *β* the DE effect sizes associated with it, and *Z* represents the covariates to be adjusted for, for example the spatially lagged expression vectors *l*_1_*, . . . , l_p_*, with *α* the coefficients associated with it. The random component *b* is assumed to have mean zero and covariance matrix Σ, which describes the correlation between cells. This correlation structure determines how the spatial correlation between cells is modeled, which is what differentiates various models: The GP models the distance between cells in a continuous way, while the BYM2 considers the adjacency of cells (Methods). An important aspect of spatial single cell transcriptomics data is the large number of cells assayed, often reaching hundreds of thousands or even millions of cells per subject. Modeling the correlation between these cells thus requires dealing with a covariance matrix of extremely large size, which introduces computational challenges. Additionally, the DE analysis usually involves multiple genes, which, even when modeled independently, further adds to the computational complexity. Therefore, alternative (approximate) approaches are necessary to address these challenges. Here we present two such approximations, one that relies on Bayesian modeling and another that models the correlation between groups of cells instead of individual cells.

The first approximation is the use of Integrated Nested Laplace Approximations (INLA) [28], within a Bayesian framework. INLA in particular uses several approximations to expedite estimation, assuming a unimodal and symmetric posterior distribution using the Laplace approximation. This speeds up computation and allows for efficient sampling. Additionally, these approximations are combined with a stochastic partial differential equation (SPDE) approach [29], which discretizes the spatial domain into a mesh, representing spatial correlation in a discrete manner. This discretization allows the use of sparse matrices, enhancing computational efficiency. A challenge of INLA is that analysts must specify suitable priors, which can influence results. However, it is highly efficient and in general yields equivalent results as classical (frequentist) inference.

The second approach consists of modeling spatial correlation not between single cells, but between clusters of cells defined by spatially partitioning the tissues (Figure 4a). By defining random effects for each cluster we model the covariance structure between clusters, instead of cells, assuming that cells within each cluster are similar to each other. We will consider three possible structures, one that assumes clusters are independent, one that considers the adjacency of clusters (BYM2) and another that models the correlation between clusters as a function of their distances (GP). By carefully selecting the appropriate model, we can accurately account for the spatial patterns. However, modeling the correlation between clusters comes with the challenge of determining the number of clusters, as there is a trade-off between processing time and how granularly we account for spatial correlation. This choice depends on the model being considered, as each model accounts for different levels of correlation depending on the number of clusters. The independent case considers only the similarity of cells within the cluster; thus if the number of clusters increases, we approach the case of not modeling any spatial correlation (Supp Fig 5). The BYM2 model considers the neighborhood of clusters; that is, it smooths the data locally. Therefore, it entails a tradeoff between insufficient granularity if few large clusters are used and loss of longer-range spatial patterns if many small clusters are used. Finally, the GP approach is able to model long ranges of spatial patterns, at the cost of not handling sparse matrices; as the number of clusters increases, the computational burden of this approach also increases.

**Figure 4:**
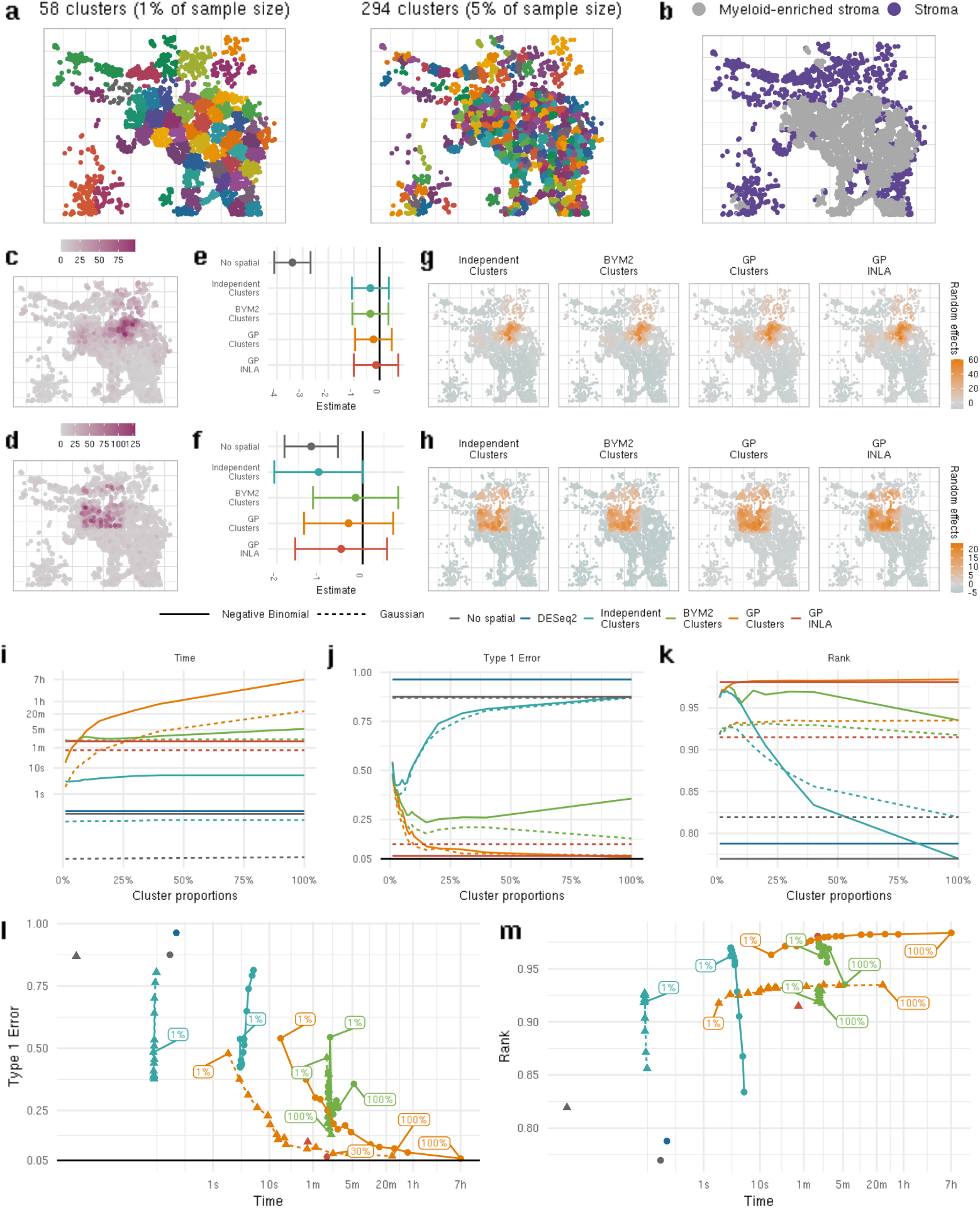
Example of clusters based on locations, each color indicates a different cluster (**a**). Representation of Stroma and Myeloid-enriched stroma niches (**b**). Visualization of results for two examples of simulations of expression of macrophages cell on stroma and myeloid enriched stroma(**c-h**), one with spatial variation generated with LGCP (**c**,**e** and **g**) and one generated with sharp boundaries (**d**,**f** and **h**). Both examples were generated in a single simulation each under the null. They assumed a Gaussian distribution, and all, except no spatial and GP-INLA, assumed 5%n clusters. Figures **c** and **d** shows the rates of gene expression, that is, the transcript count over total transcript count for each cell. Figure **e** and **f** show the estimate of difference of expression between stroma and myeloid enriched stroma with corresponding confidence/credible intervals. Figures **g** and **h** show random effects from a single model run. Figures **i-l** show results obtained based on results from repeated simulations. **i** shows average processing time (in minutes), with the y axis bounded at 50 minutes for better visualization. **j** shows Type 1 error obtained based on 600 repetitions, **k** shows Spearman correlation between estimates and true coefficients for simulated data. Values on the x-axis are the proportion of the data considered in each cluster.

#### Simulation to analyze spatial variation impact on type 1 error and gene ranking

The different models and numbers of clusters were benchmarked in a simulation study, where we compared models without any spatial component (No Spatial model and DEseq2), a model with independent clusters, the BYM2 and GP models. We also compared the GP model with INLA approximaiton, which is obtained only for the case where the correlation between all cells is modeled. The number of clusters is presented as a percentage of the total number of cells (%*n*) so that recommendations can be made for general datasets (Figure 4a). Data was generated based on a real dataset on FFPE non-small-cell lung cancer tissue samples. Here, we compared 5878 macrophages in the stroma and myeloid enriched stroma niches (Figure 4b). We start by looking at two data examples; in the first example, gene expression counts were generated based on a log Gaussian Cox process (LGCP) [30]; in the second, an artificial spatial pattern of a square was used. In both cases (Figures 4c-h), the gene expression was generated under the null hypothesis that the expression does not differ by niche. Then, in the LGCP case, we performed several simulations under the null and alternative hypotheses (see Methods for more details) to compare different models and cluster number based on processing time, Type 1 Error, Power and correlation between ranking of genes based on true and estimated coefficients. Both Negative Binomial and Gaussian distributions were considered.

Figure 4c shows that, due to the spatial pattern in the first setting, cells in the upper right part exhibit the highest expression rates even under the null hypothesis, which is a location that mostly contains cells in the myeloid-enriched stroma niche (Figure 4b). For the second setting, Figure 4d shows that cells have higher expression inside the artificial square generated. In both simulations (Figures 4e and 4f), not accounting for spatial variation would identify the gene as differentially expressed between niches. When accounting for this correlation, regardless of the model, we achieve the correct conclusion that there is no difference between the niches. In Figures 4g-h we can further investigate the spatial random effects of each model. We see that the pattern obtained by the independent cluster, BYM2 and GP are similar, and all models are able to identify the patterns generated by the spatial effects.

GP models have the highest computational times, taking up to 400 minutes with less than 6000 cells, assuming a Negative Binomial distribution. Shorter processing times can be obtained by modeling the correlation of clusters, instead of individual cells, or assuming a Gaussian distribution (Figure 4i). Using Laplace approximations, such as the BYM2 or GP-INLA, reduces the average processing time by more than 90% compared to the GP model without using clusters. Neglecting to account for spatial correlation leads to an uncontrolled Type 1 error greater than 80% (Figure 4j). Therefore, any of the spatial models, even with a small number of clusters, yields better control of false positives. GP models demonstrates the most effective control of type 1 error. When assuming a Gaussian distribution, the Bayesian version of GP leads to inflated type 1 error, whereas the frequentist approach (GP clusters) show similar results as when assuming a Negative Binomial distribution. Notably, it is possible to obtain good control of false positives even when modelling correlation of clusters with number corresponding to 25%*n*. With this cutoff, the processing time and type 1 error of GP-clusters and GP INLA are similar (Figure 4l). Models that perform smoothing of the spatial correlation, such as BYM2, fail to fully account for the spatial confounding. This leads to uncontrolled Type 1 error; however BYM2 is still able to provide good correlation between ranks of true and estimated coefficients (Figure 4k), which can be useful for screening top associated genes. Once again, GP models provide the best ranking of differentially expressed genes, even when assuming a small number of clusters. But unlike the Type 1 error, for spatial models, better ranking of genes is obtained when assuming a Negative Binomial distribution, compared to Gaussian.

To summarize, it is important to account for the spatial variability of gene expression to better control the Type 1 error and correctly rank the genes. Doing so relies on modeling the correlation between all pairs of cells, which may not scale to large number of cells and may lead to long computational time, which is infeasible in most applications. To achieve acceptable computation times, we presented alternative approaches based on approximations, including modeling the correlation between clusters, instead of individual cells and using a Laplace approximation in a Bayesian framework. By considering clusters, we can significantly decrease the computational time, and obtain results very close to those obtained from full modeling of correlations, especially for larger numbers of clusters, when using a GP or BYM2 model. Finally, using the Laplace approximation of INLA for the GP model allows modeling the individual level data with shorter processing time.

### 2.4 Analysis of CosMx NSCLC dataset

To further benchmark the different methods and provide recommendations on suitable analysis approaches, we applied the methods discussed in Sec 2.3 to the spatial CosMx NSCLC transcriptomics dataset previously explained. The analysis aims to compare expression of 980 genes on 14740 tumor cells in tumor interior and tumor stroma boundary (Figure 5a). Following the recommendations of Sec **??**, overlap ratio metrics were obtained, and 645 genes with overlap ratio metric less than 1 were considered for the analysis. We compare models with different adjustments for spatial correlation, while accounting for unobserved transcript misassignment from segmentation error and assuming a Gaussian distribution. The number of clusters for each model were chosen following the results in Figure 4: For the independent cluster model, we chose 5% of the total number of cells, which corresponds to 737 clusters, based on results for Type 1 error and ranking. For BYM2 and GP models, we chose 25% of 14740 cells, corresponding to 3685 clusters, as it leads to faster computational time, while maintaining performance similar to modeling with individual cells.

**Figure 5:**
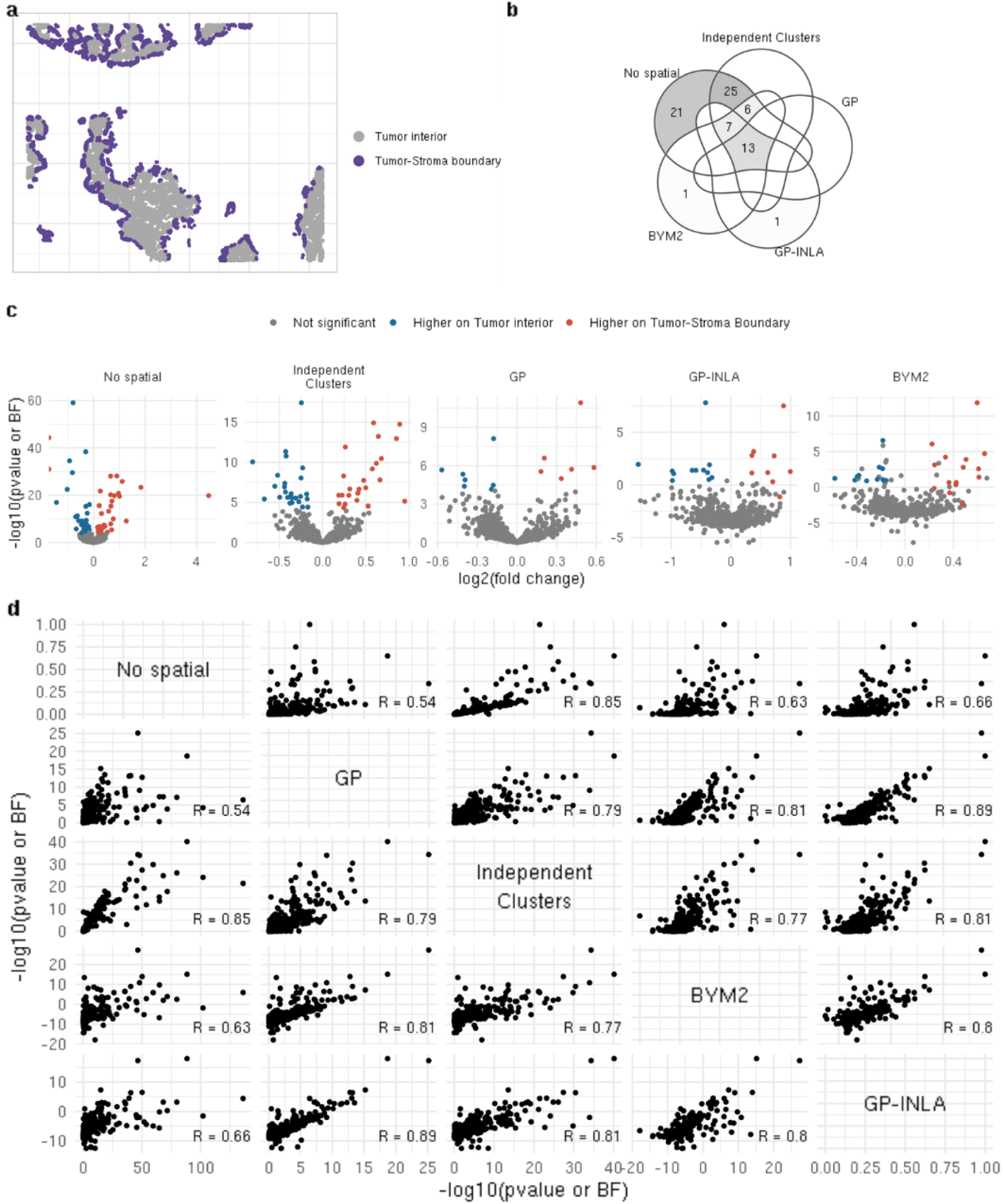
**a**, locations of 14740 cancer cells on tumor interior and tumor-stroma boundary. **b**, venn diagram comparing significant genes identified in common by each model. **c**, volcano plots with estimates for different models and -log10(pvalue) for no spatial, independent clusters and GP models, and -log(Bayes Factor) for BYM2 and GP-INLA models, under Gaussian distribution. **d**, scatter plot comparing p-values or bayes factor of each model. For model without spatial modeling, independent cluster and GP genes with p-values lower than 0.05/980 were considered significant, whereas for BYM2 and GP-INLA significance was determined based on (1-0.05/980)*100% credible interval.

Figure 5c shows volcano plots with up or down regulated genes using each model. Since the BYM2 and GP-INLA models are obtained under a Bayesian framework, Bayes Factors (BF) are used in place of p-values. However, BF was only used for visualization and ranking of genes and significant DE genes were determined based on credible intervals. This is due to the fact that, based on our simulations, the BF can be sensitive to the choice of prior considered, whereas the estimates and credible intervals are more robust (Supp Fig 6).

Starting with the naïve model that does not account for spatial correlation, we see greater significance compared with other approaches, with − log 10 p-values around 60. Also, from Table 1 we see that more genes were deemed significant compared with other methods. These findings agree with the simulation results showing higher Type 1 error for this approach. From the 72 genes found significant at a 0.05 significance level after Bonferroni adjustment, only 13 were also identified by the GP model.

**Table 1:** Number of down/up regulated or unchanged genes under each model.

| Model | Down regulated | Unchanged | Up regulated |
| --- | --- | --- | --- |
| No spatial | 35 | 573 | 37 |
| Independent clusters | 26 | 594 | 25 |
| GP | 7 | 632 | 6 |
| GP-INLA | 12 | 624 | 9 |
| BYM2 | 12 | 618 | 15 |

For models that account for the spatial correlation, we compare the number of significant DE genes (Table 1), the correlation between p-values (Figure 5d) and the correlation between fold change estimates (Supp Fig 7). The results from the independent cluster method are closest to the No Spatial method, followed by BYM2 then GP. The GP models are better able to account for the spatial correlation between cells by selecting less potential false positives and better controlling the Type 1 error; this mirrors the simulation results. The independent clusters and BYM2 models select less significant genes than the No Spatial model, which might show better control of false positives. The two GP models—the one under the frequentist framework that models the correlation between clusters of cells, and the one under a Bayesian framework with Laplace approximations that models correlation between individual cells—have very similar results: The correlation between p-values and Bayes Factor (BF) is 0.941, the correlation between estimates is 0.948, and the two methods select essentially the same DE genes (Figure 5b). The high correlation between p-values and BF values is notable given the potential sensitivity of BF to the choice of priors. This finding further supports the use of BF to rank significant genes.

To demonstrate the importance of accounting for the spatial correlation between cells, we look at four genes as specific examples: HLA-A, S100A6, TYK2 and CXCL5 (Figure 6). HLA-A demonstrates consistent p-values and fold change estimates between the No Spatial model and the GP models, under a frequentist approach. Even though the residuals from the No Spatial model have a spatial pattern, they are uncorrelated with the niche; therefore, the correlations did not influence p-values and fold changes.

**Figure 6:**
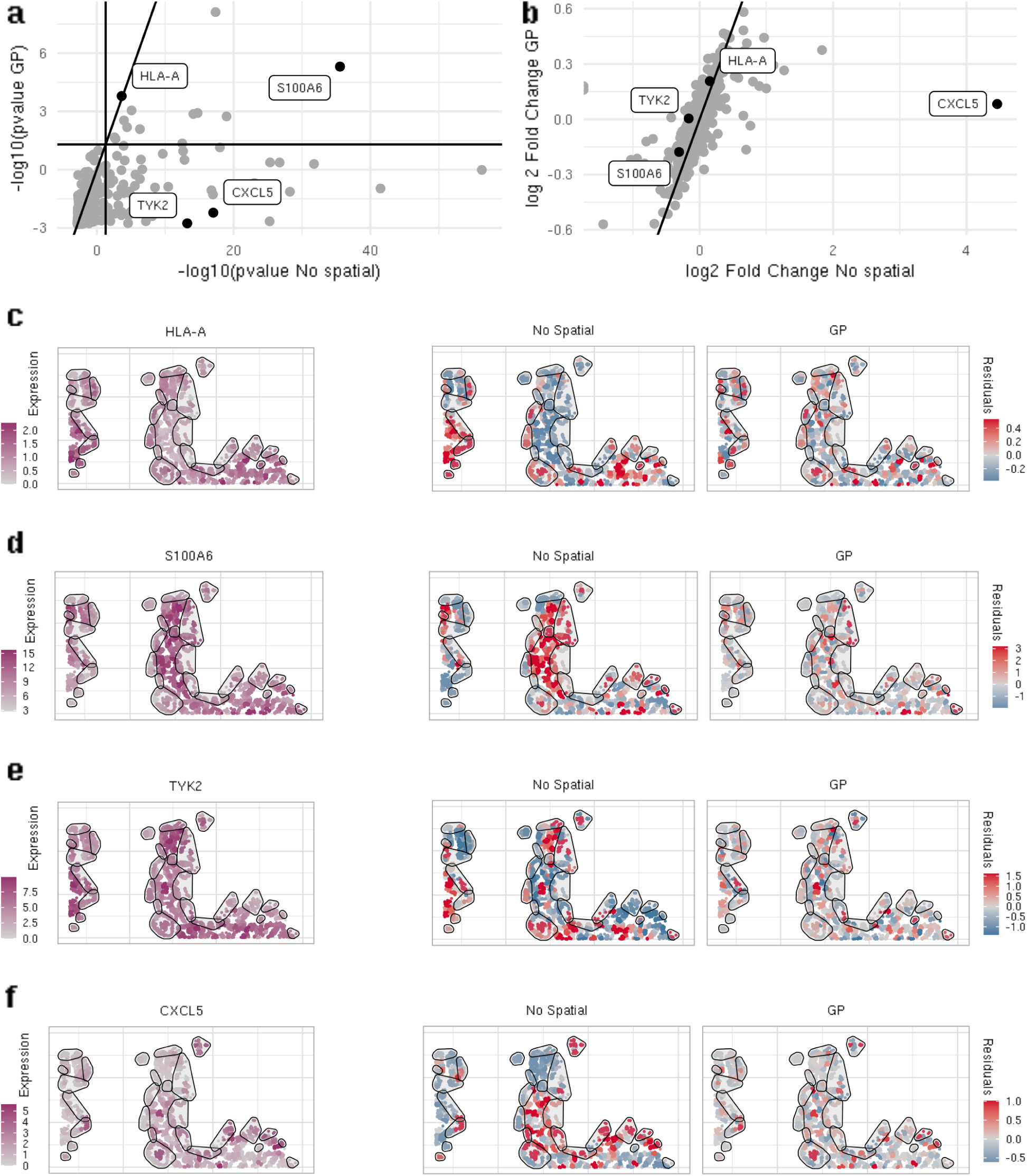
Comparing results obtained between No Spatial model and Gaussian Process. P-values and Fold change estimates are shown in Figures a and b respectively. Figures c-f show normalized expression and residuals obtained under no spatial and GP for specific genes. For the normalized counts and residuals, cells within the same niche were grouped and the average value is shown, to allow for a better visualization of the patterns with data less noisy.

For S100A6, both models resulted in similar estimates, but the GP model showed a lower p-value compared to the No Spatial model. When examining the residuals from the No Spatial model, we note a pattern associated with the tumor interior and tumor interior boundary. This pattern likely contributes to the decreased p-value observed in the GP model, since the random effects might model parts of the niche effect. However, in both cases we still have evidence of higher expression of this gene in the tumor interior.

In contrast to the above findings, the No Spatial model identified TYK2 as significant, whereas this gene was not found significant using the GP model. Under the No Spatial model, TYK2 exhibited a spatial pattern in both expression and residuals, with pattern that differ from those observed in S100A6. Unlike S100A6, the spatial patterns are not different between the niches. When accounting for spatial correlation, the gene is no longer differently expressed in tumor interior and tumor stroma boundary.

Finally, CXCL5 was identified as significant by the No Spatial model with a fold change greater than 20, while the GP model showed a fold change close to 1, rendering it non-significant. This gene clearly has a spatial expression pattern, with a few small clusters exhibiting much higher average expression, particularly in the tumor interior niche. This correlation leads the No Spatial model to overestimate the expression on the boundary relative to the interior. In contrast, by accounting for spatial correlation, the GP model reveals no difference in expression between the two niches. The expression pattern of this gene highlights the impact of spatial correlation on obtaining reliable DE analysis.

### 2.5 Analysis of FFPE Colon Cancer Tissue

Using a publicly released CosMx 6,000-plex colon cancer dataset [31] with several tertiary lymphoid structures, we explored how B cells responded to the number of immune cells within a 15 micron radius (Figure 7a). Among other functions in tumors, B cells form tertiary lymphoid structures (TLS), consisting of B cell dense germinal centers surrounded by T cells. TLS’s are often associated with positive cancer outcomes [32]. The number of immune cells (including other B cells) in a B cell’s immediate neighborhood is a continuous covariate which effectively serves as a measurement of how close that B cell is to the center of a TLS, and may provide insight into what genes are differentially regulated in these clinically relevant functional regions.

**Figure 7:**
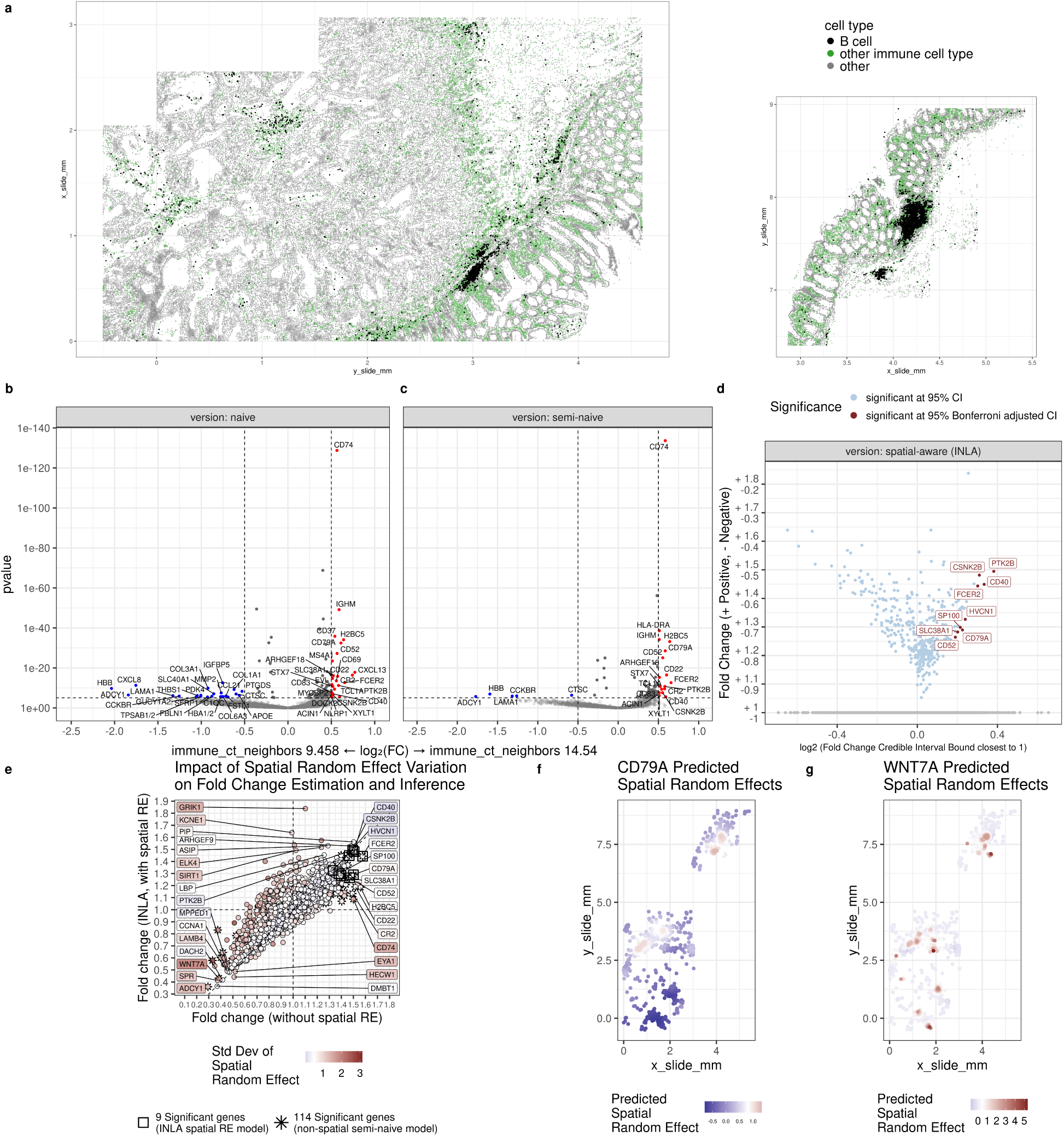
**(a)** Colon tissue colored by cell type category; B cells (black), other immune cell types (green) include macrophage, mast, mDC, monocyte, neutrophil, NK, pDC, plasmablast, and T cell. ‘Other’ category includes epithelial and stromal cell types. Volcano plots for **(b)** naive model without gene filtering or covariate adjustment, **(c)** semi-naive model, with gene filtering, covariate adjustment, but without spatially correlated random effect (SRE), and **(d)** spatial-aware model results which include gene filtering, covariate adjustment, and SREs. For the bayesian GP-INLA model shown in **(d)**, the x-axis coordinate represents the fold change (FC) credible interval (CI) bound closest to the null hypothesis (1), on log2-scale. FC distance is plotted on the y-axis for all genes with 95% CI’s which excluded 1. Genes with *α*=(1-0.025/2922)% FC CI’s which excluded 1 are highlighted in red. **(e)** FC estimates from semi-naive model without SREs (x-axis) plotted against FC estimates from spatial-aware SRE model (y-axis), colored by estimated standard deviation of SREs. Genes with higher variability in SRE (red color), are more likely to lose significance in spatial-aware model. Predicted SRE for **(f)** CD79A and **(g)** WNT7A are examples of low and high variability in SRE. WNT7A becomes not-significant in the spatial-aware model, while CD79A remains significant in presence of SRE.

To compare the tradeoffs of different DE analysis workflows and their impact on inference, we considered three different models employing methodology described thus far, with results highlighted in the volcano plots (Figure 7b-d). For the first model (naive) (Figure 7b), we modeled all genes; neither filtering to account for segmentation error, nor adjusting for expression of the gene in neighboring cells of other cell types as described in (2.2). For the second model (semi-naive) (Figure 7c), we removed genes from analysis with overlap ratios greater than 1, and used neighbor cell expression as a control variable as described in (2.2). For the third model (spatially aware) (Figure 7d), we included a spatial random effect using the GP-INLA approach described in (2.3), in addition to the gene filtering and covariate adjustment methods used in the semi-naive model. All models were fit using negative binomial regression.

The naive model discovered 141 significant genes based on a bonferroni p-value threshold of *p <* 0.05*/*5914, composed of 35 down-regulated genes (for which the expression increases when a B cell is surrounded by fewer immune cell neighbors), and 106 up-regulated genes (for which expression increases when surrounded by a larger number of immune cells). Unsurprisingly, the naive analysis highlighted a number of genes which a biologist may deem implausible, particularly among down-regulated genes not predicted to be expressed in immune cells (for example, ADCY1, COL3A1)[23, 22], whose expression would challenge the cell type identity of “B cell”, and whose significance indicates likely confounding between the primary covariate (number of immune cell neighbors) and misassigned transcripts arising from segmentation error. The semi-naive model automatically removed 3,214 genes from analysis by filtering genes with overlap ratios *>* 1, and discovered 114 significant genes based on a bonferroni p-value threshold *p <* 0.05*/*2700, 98 of which were up-regulated DE genes. All genes which were significant in the semi-naive model were also significant in the original naive model. Finally, the spatially aware model, which included a spatially varying random effect, discovered 9 significant genes, all of which are up-regulated in the presence of a larger number of immune cell neighbors. For the bayesian spatial-aware model INLA, which does not compute p-values, we determined genes as signficant if their *α* = 1 − 0.025*/*2700 level credible interval for fold change excluded the value of 1. The credible interval can correspond to an acceptance region for a bayesian hypothesis test under assumptions described in [33] and [34]. This smaller set of genes represents a high-confidence gene set composed primarily of genes which can be defended based on existing literature. For example, FCER2 is known to initiate IgE-dependent antigen uptake and presentation to T cells [35, 23], which is plausible given the proximity of B cells to T cells in high-immune-cell TLS regions. Another DE gene, SP100, is a tumor suppressor gene and thought to play a roles in tumorigenesis, immunity, and gene regulation [36, 23]. In summary, of the 141 genes found significant by the naive analysis, 27 can be explained as resulting from errors in cell segmentation, and 105 could be explained by spatial variability unrelated to the biological question. The methods proposed here thus identified 93.6 percent of the naive method’s findings as potentially false discoveries.

To give insight into the impact of the spatial random effect on inference, we plotted the fold change estimates for each gene from the semi-naive model against the fold change estimates from the spatially aware model (Fig 7e), and colored the points by the estimated standard deviation of the spatial random effects from the spatially aware model. Genes with higher variation in spatial random effects tend to be less likely to be significant in a spatially aware model, as variability in the primary covariate can be increasingly explained by a spatially smoothed random effect. To further aid interpretation, we plotted the predicted random effects for CD79A (Fig 7f) and WNT7A (Fig 7g). WNT7A had high spatial variation, and was significant in the semi-naive model, but not in the spatially aware model. We can see based on the predicted random effects that there are apparent hyper-specific hot-spots of WNT7A expression that explain away the variation in expression we might otherwise attribute to our primary DE covariate. CD79A remained significant in the spatially aware model, and has comparably lower variation attributable to the spatial random effect, with the variation being visually smoother compared to the random effects for WNT7A. For this analysis of 2,922 B cells, average model estimation time per gene was 0.11 seconds for the naive model, 0.13 seconds for the semi-naive model, and 21.6 seconds for the spatial-aware model.

## 3 Discussion

Spatial transcriptomics data presents several analytical challenges, some of which overlap with issues encountered in single-cell studies, such as addressing count data with sparse signals; however, there are also distinct challenges specific to spatial transcriptomics, including managing the impact of cell segmentation errors and accounting for spatial correlations. In this article, we explored various modeling approaches focused on this data type, considering their computational efficiency, a crucial concern given the scale of spatial transcriptomics datasets. To facilitate their adoption, the methods discussed in the paper, including the new GP cluster method, have been implemented in an R package, smiDE, which will be made available on GitHub along with extensive documentation.

We demonstrate that while the Negative Binomial distribution is ideally suited for over-dispersed count data, Gaussian regression models can be viable alternatives with suitable normalization. Notably, the Gaussian distribution offers accurate approximation, usually with shorter run time, especially when accounting for the spatial correlation. This is particularly valuable in large datasets, where faster methods can enhance efficiency without compromising accuracy.

Addressing misassigned transcripts arising from cell segmentation is crucial. We introduce a data-driven metric for flagging and removing genes from a DE analysis which are likely to be driven by imperfect segmentation. Additionally, we propose a useful proxy covariate that can be easily calculated and integrated into DE regression models. We show that covariate adjustment and gene filtering both reduce false positive discoveries arising from segmentation error in DE analysis.

Furthermore, we emphasize the critical importance of accounting for spatial correlation to mitigate the risk of inflated Type 1 errors, as already known [7]. We further propose a new variant of mixed effect models that considers correlation between clusters, which deals with spatial correlation while improving computational time. Based on comprehensive simulations, we observe that the No Spatial model and DESeq2, although exceptionally fast, fail to control the Type 1 error and fail to provide accurate rankings of top genes.

Focusing on the ranking of genes is of particular interest in exploratory analyses, in which the goal is to find the most differently expressed genes, and to generate hypotheses for further studies. In this context, the Gaussian Process (GP) cluster model with a Negative Binomial distribution stands out—it provides reliable rankings even with as few as 10%*n* clusters. When computation becomes more challenging due to large number of cells and/or genes, the independent cluster model with 5%*n* clusters is a viable computational alternative—it maintains good rankings while considerably reducing runtime.

For confirmatory analyses, accounting for Type 1 error is critical, and simulations indicate that the GP model is the most robust approach. This prompts a decision between frequentist and Bayesian inference. While Bayesian inference expedites computational time through approximations, it introduces nuances related to the choice of priors, which can impact the stability of results. Furthermore, Bayesian methods provide inference through credible intervals but do not provide traditional p-values, relying instead on Bayes Factors. Opting for frequentist inference ensures stable and robust results across various scenarios but demands more computational time, nearly double that of Bayesian approximations. In this scenario, considering a Gaussian distribution and 25%*n* clusters can balance processing time and model accuracy (Figure 4l). In our simulations, the frequentist GP model with 25%*n* clusters, and the Bayesian approach lead to similar results; however, processing time of GP-clusters increases at a higher rate than GP-INLA. Therefore, for very large number of cells, GP-INLA might be preferable.

Our analysis focused on two-niche comparisons. However, extending to multiple comparisons, whether pairwise or one-vs-all, is possible. This extension does not significantly impact the computational time of GP clusters. However, it could impact the computational time of calculating Bayes Factors when using GP-INLA. Therefore, the number of relevant niches in the analysis should be considered when selecting the preferred model. While computational time currently poses a challenge, there are emerging methods to expedite analyses, such as NEBULA [15], which can be expanded to allow for modelling spatial variation. Therefore, it may soon be unnecessary to choose between speed and accuracy.

When dealing with spatial statistics, a key concern is spatial confounding, a well-recognized challenge in the field [37, 38, 39, 40]. Spatial confounding occurs when the spatial variation closely aligns with the covariate of interest, such as tissue niche. In such scenarios, attempting to adjust for spatial variation can diminish the informative signal associated with the niche, which can result in reduced statistical power and complications in finding meaningful associations. If the covariate is able to entirely account for the variability in expression, then there will be no loss in power; see simulation results in Supp Fig 8. Although power is not directly comparable across models due to inflated Type 1 errors, our simulations find that modelling spatial correlation improves power compared to the “No Spatial” model and DEseq2). However, if the niche fails to explain all the variability and is multicollinear with the spatial random effect, power may be compromised. The field has not reached a consensus on the optimal approach to address spatial confounding [41], and it is a limitation present in most spatial analyses that must be kept in mind. On the other hand, as demonstrated through our analyses, ignoring spatial correlation can lead to inflated Type 1 errors and abundant false discoveries. In practice, some genes found significant by only “No Spatial” models may still be of interest, although they deserve much greater skepticism than their p-values suggest.

Our discussion of spatial modeling primarily focused on individual samples and not on comparing between samples. For within-sample questions, we recommend performing differential expression separately for each sample. This approach reveals not only each gene’s average effect size, but the diversity of effects across samples. Results for each gene can then be summarized using meta-analysis techniques such as inverse variance weighting [30]. This approach of stratifying by sample is usually feasible, as most studies have large numbers of cells per sample and relatively few samples.

## 4 Methods

### 4.1 Generalized Linear Mixed Models

Assume we have *n* cells and *p* genes, let *Y_ig_* represent the expression of the *g*th gene in *i*th cell, with conditional expectation *µ_ig_* modeled as

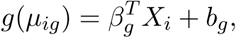

where *X* is the design matrix for fixed effects, with coefficients *β*, and the random effects *b* follow a Gaussian distribution with covariance matrix Σ(*θ*) that models the spatial correlation between cells. This correlation can be modeled in several ways. The first corresponds to a Gaussian Process (GP), which assumes an exponential covariance model Σ(*θ*)*_ij_* = *θ* exp(−*ρd_ij_*) where *d_ij_* is the distance between cells *i* and *j*, obtained based on their locations and *ρ* is a scale parameter.

Alternatively, we can model the data using the BYM2 model [27], with the correlation matrix Σ(*θ*) = *θ*((1 − *ϕ*)*I* − *ϕQ*^−1^)^−1^, where *Q* represents a precision matrix obtained -based on the adjacency matrix of cells and *ϕ* is the proportion of variability of the random effects due to spatial components. This model assumes that the expression for the *i*th cell depends on the expression of its neighbors.

Inference about (*β, θ*) can be performed under two frameworks: using Maximum Likelihood or with Laplace approximations (INLA). One challenge we might face is dealing with Σ*_n_*_×_*_n_* when the number of cells is large, since the matrix size increases rapidly, slowing down the estimation. While the BYM2 model handles sparse matrices for faster computation, this may not be efficient for the GP model. To address this issue, INLA uses an approximation that discretizes the space using triangularizations, which approximates Σ(*θ*) by a sparse adjacency matrix for faster computation.

Here we propose another approximation, which consists of modeling the correlation between clusters of cells instead of each individual cell, which can also improve computation speed, while maintaining similar conclusions.

### 4.2 Random Effects for Clusters

We consider clustering techniques, such as K-means, to group cells that are close to each other based on their locations. Each cluster is then associated with a random effect. Under a GP model, *d_ik_* measures the distance between centers of clusters *i* and *k*. If the BYM2 model is used, we consider the adjacency matrix between clusters. Alternatively, we may assume that cells within a cluster share the same random effect, but clusters are independent of each other. Using this approach we obtain a *K* × *K* matrix, where *K* is the number of clusters. The number of clusters is a tuning parameter that determines the amount of spatial correlation in the model.

### 4.3 Simulations

We used simulation studies to compare different spatial models and better understand the impact of the number of clusters. One approach to simulate data considering the real spatial correlation is by using a Log Gaussian Cox Process (LGCP). For this, we assume the expression of the gene for the *i*th cell, *Y_i_*, follows a Poisson distribution with mean *µ_i_*. Specifically,

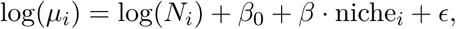

where *N_i_* is the total transcripts counts for the cell, *β*_0_ is an intercept to guarantee that cell counts are on the scale similar to real data, niche*_i_* indicates the niche of cell *i*, with the corresponding coefficient *β*. We further assume *ϵ* follows a Gaussian distribution with mean 0 and exponential covariance matrix. The covariance matrix is obtained based on the distances of cells in a real data example.

Type 1 error was evaluated by simulating data with *β* equal to zero, for 600 iterations. To analyze the ranking of the models, 100 simulations were performed, by generating 21 genes, 6 of which with null effect, and others with effects varying from 0.1 to 1.5. Then, Spearman correlation between estimated and true *β*’s as well as p-values were calculated to verify if the models were able to correctly rank genes by their associations.

### 4.4 An Overlapping Cells Ratio Metric for Cell Type Specific Gene Filtering

We develop an Overlapping cells ratio metric for filtering out the most implausible genes in DE which are most likely to arise from segmentation errors.

In a DE model for a specific cell type, let *t* denote the cell type of interest and *t*^′^ denote all other cell types. Intuitively, when analyzing a specific cell type, we expect that a gene may be prone to error from overlapping segmentation if it has higher expression in spatially close neighboring cells of other cell types *t*^′^, than the cell type of interest, *t*.

With *n_t_* denoting the number of cells of type *t* and *n_t_′* denoting the number of cells in all other cell types *t*^′^, let *W* (*r*) denote an *n_t_* × *n_t_* spatial neighborhood matrix, where each element *w_ij_* = 1 if cell *j* is within distance *r* of cell *i*, and 0 otherwise. Let *M* (*r*) denote a *n_t_′* × *n_t_′* diagonal matrix with diagonal elements 1*/m_ii_* indicating the number of neighbors to cell *i* within radius *r* in other cell types 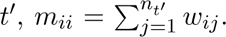 Further let *y*(*t*)*_g_* and *y*(*t*^′^)*_g_* denote the *n_t_* and *n_t_′* length vectors of expression for gene *g*. Then, the average expression of the gene in neighboring cells of other cell types is given by 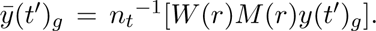 Letting 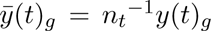 denote the average expression of gene *g* for the type *t* cells of interest, the ratio

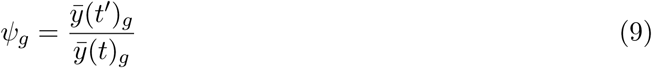

is a metric by which we quantify the susceptibility to confounding bias from overlapping or erroneous cell boundaries. Genes with *ψ_g_ >* 1 are higher expressed, on average, in neighboring cells of other types than the cell type of interest; hence the observed number of transcripts in analyzed cells may have an unacceptably high rate of false positives.

### 4.5 Analysis using CosMx NSCLC Data

#### 4.5.1 Choosing Parametric Distribution for DE regression model

Using the CosMx NSCLC dataset [1] with 487,819 total cells from 5 NSCLC tissues, we compared three parametric regression models (Negative Binomial RE, Poisson RE, or Gaussian RE) in terms of computation time, concordance in effect size estimation, and significant gene calls. Specifically, we ran DE regression models for 7 cell types (macrophage, T-reg, T-CD8, T-CD4, plasmablast, neutrophil, and tumor). We used the rank-normalized and inverse distance weighted total expression of neighboring cells of other cell types as described in Eq. (7) as an adjustment covariate in our regression model. A categorical covariate ‘niche’ annotates the functional region of tissue the cell resides in, and was treated as the primary covariate of interest in the regression models. This ‘niche’ covariate had 9 categories (tumor interior, storma, tumor-stroma boundary, myeloid-enriched stroma lymphoid structure, immune, plasmablast-enriched stroma, and macrophages). The emmeans package [42] was used to calculate ‘one vs. the rest’ contrasts for each niche, such that for each cell type, we assessed whether a gene was DE in a particular niche compared to the cell-weighted average of all other niches. This constitutes 9 contrasts per analyzed gene. We further subset the number of analyzed genes for each cell type using Eq. (8), removing genes with overlap ratio metric *ψ_g_ >* 1.

#### 4.5.2 Countermeasures for cell segmentation error

We compared the behavior of the gene flagging metric and covariate adjustment method across DE analyses of macrophage, T-CD8, T-CD4, and T-reg cell types across 5 patients in the CosMx NSCLC dataset as described above. For each cell type, we fit regression models and tested whether genes were DE in any of the 9 ‘niches’ compared to a cell-weighted average of other niches. For each cell type, we fit models both with and without a control variable. For models including the control variable, the covariate was defined as the rank normalized, inverse-distance weighted total expression of neighboring cells of other cell types as described in Eq. (7). To determine the effectiveness of the overlap ratio metric, we considered an unfiltered analysis of all genes, as well as removing those with overlap ratio metric *ψ_g_ >* 1. When assessing plausibility of significant genes based on their annotation for “Immune Cell Specificity” in Human Protein Atlas [22, 23], we considered them to be “Enhanced or Enriched” in T-reg,T-CD8,T-CD4 if the annotation contained “enhanced” or “enriched” in the description of Immune Cell Specificity, in addition to any of the phrases “T-cell”, “T-reg”, or “Treg”. For the macrophage cell type, a gene was considered to be “enhanced/enriched” if the description contained the phrase “enhanced”, “enriched”, in addition to “monocyte”, “macrophage”, “myeloid”, or “DC”. Genes were considered as “enhanced/enriched (another immune cell type)” if they contained “enhanced” or “enriched” in their description, but did not meet the cell-type specific criteria described above. Genes were labeled as “Low Immune Cell Specificity” or “Not Detected in Immune Cells” if this was the exact description for the gene provided in Human Protein Atlas.

The authors declare that they have no competing interests.

## Supplementary Material

**Supp Fig 1:**
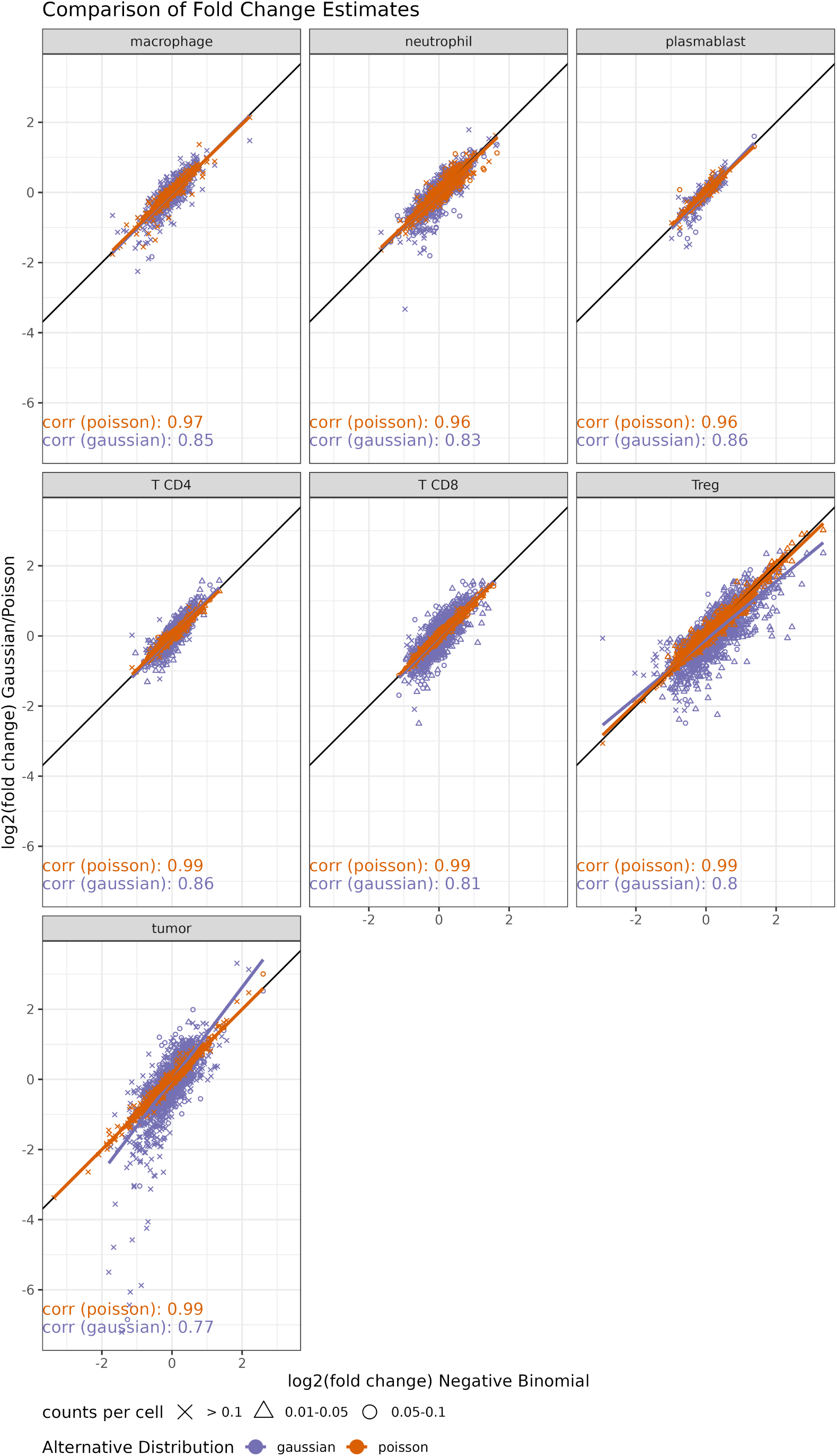
Comparison between Effect Size estimates by Family.

**Supp Fig 2:**
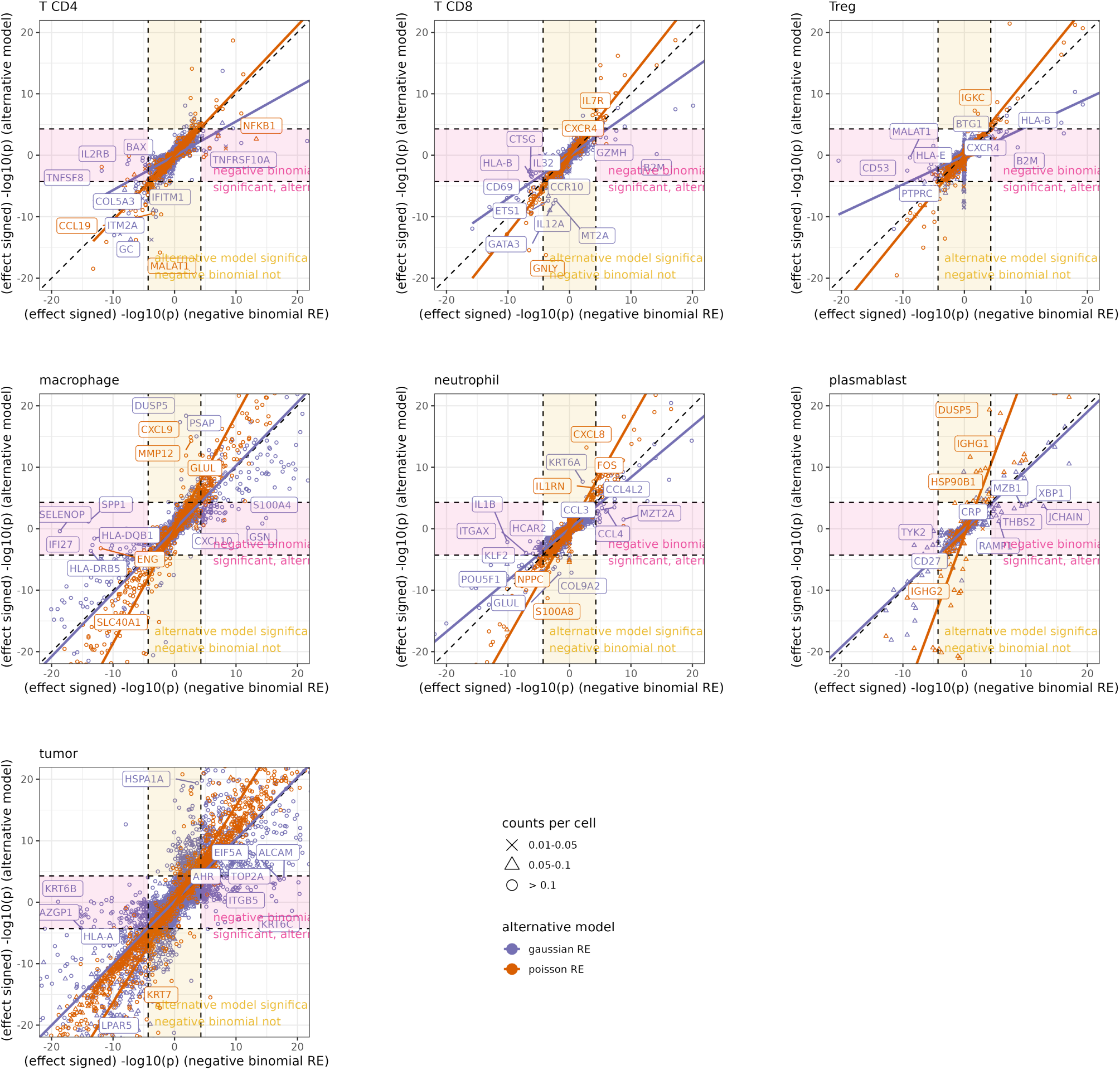
Comparison in signed log10(p) by model type.

**Supp Fig 3:**
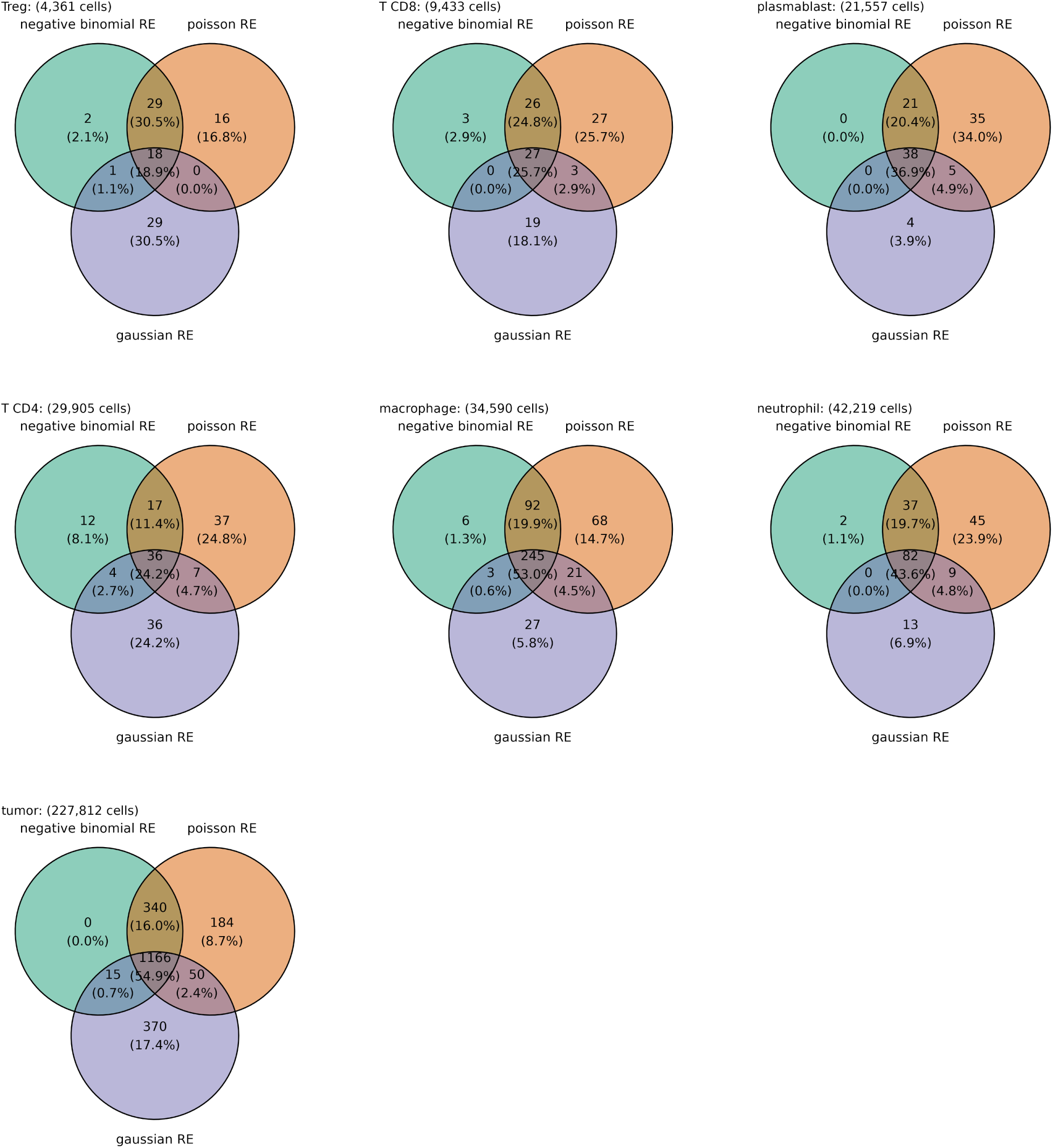
Shared significant genes by model type.

**Supp Fig 4:**
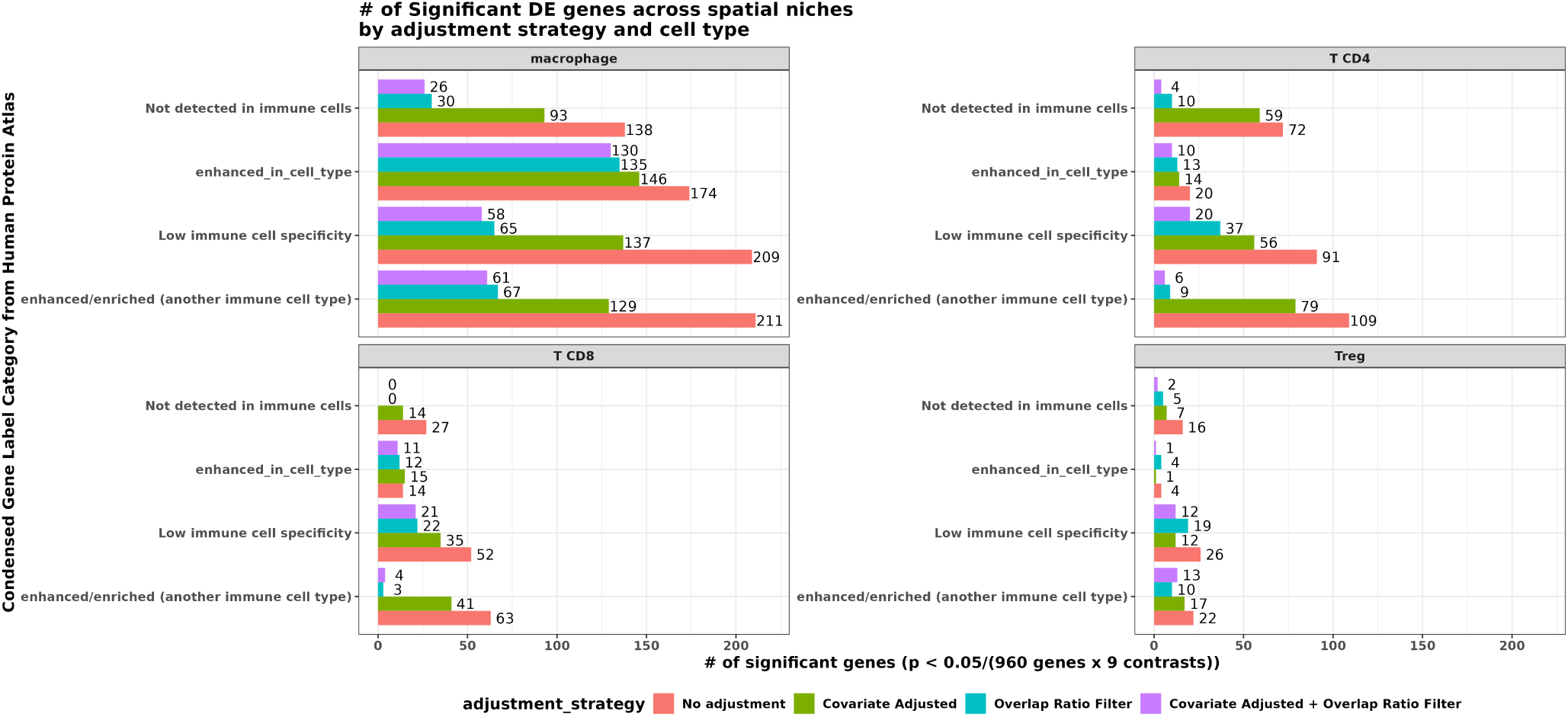
Number of significant genes using different strategies to adjust for transcript misassignment from segmentation error.

**Supp Fig 5:**
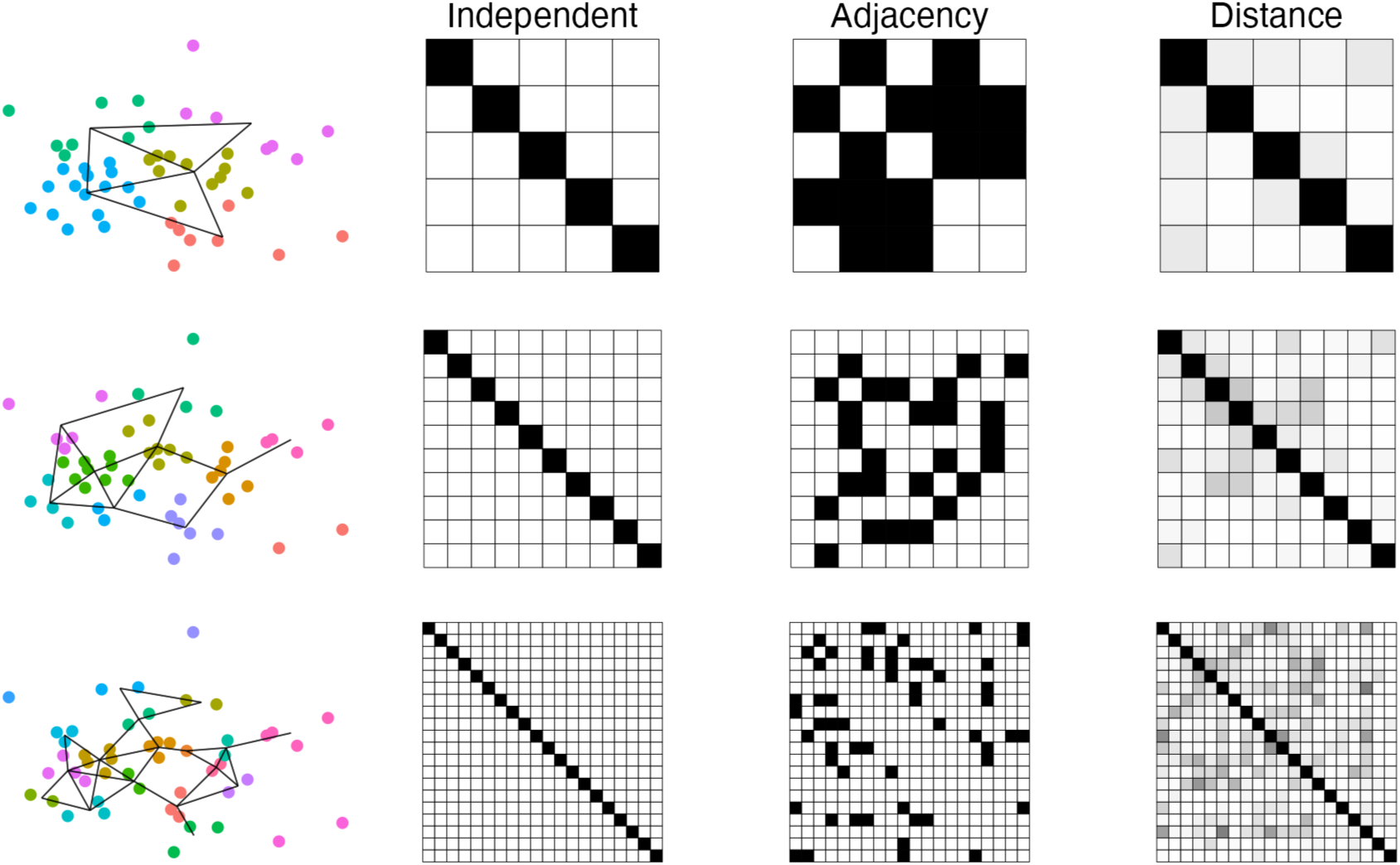
Representation of covariance matrix in each model for different number of clusters. Simulation of 50 cells and clusters of size 5, 10 or 20, represented by different colors. Lines between cells indicates if clusters are adjacent. For each cluster we have a covariance matrix showing association between them, with values from 0 (white) to 1 (black). For the independent case we have a diagonal of 1 and off diagonals of zeros. For adjacency matrix we have entries of 1 for clusters that are adjacent to each other and 0 otherwise. For distance matrix we have values between 0 and 1, with bigger values indicating shorter distance between clusters centroids.

**Supp Fig 6:**
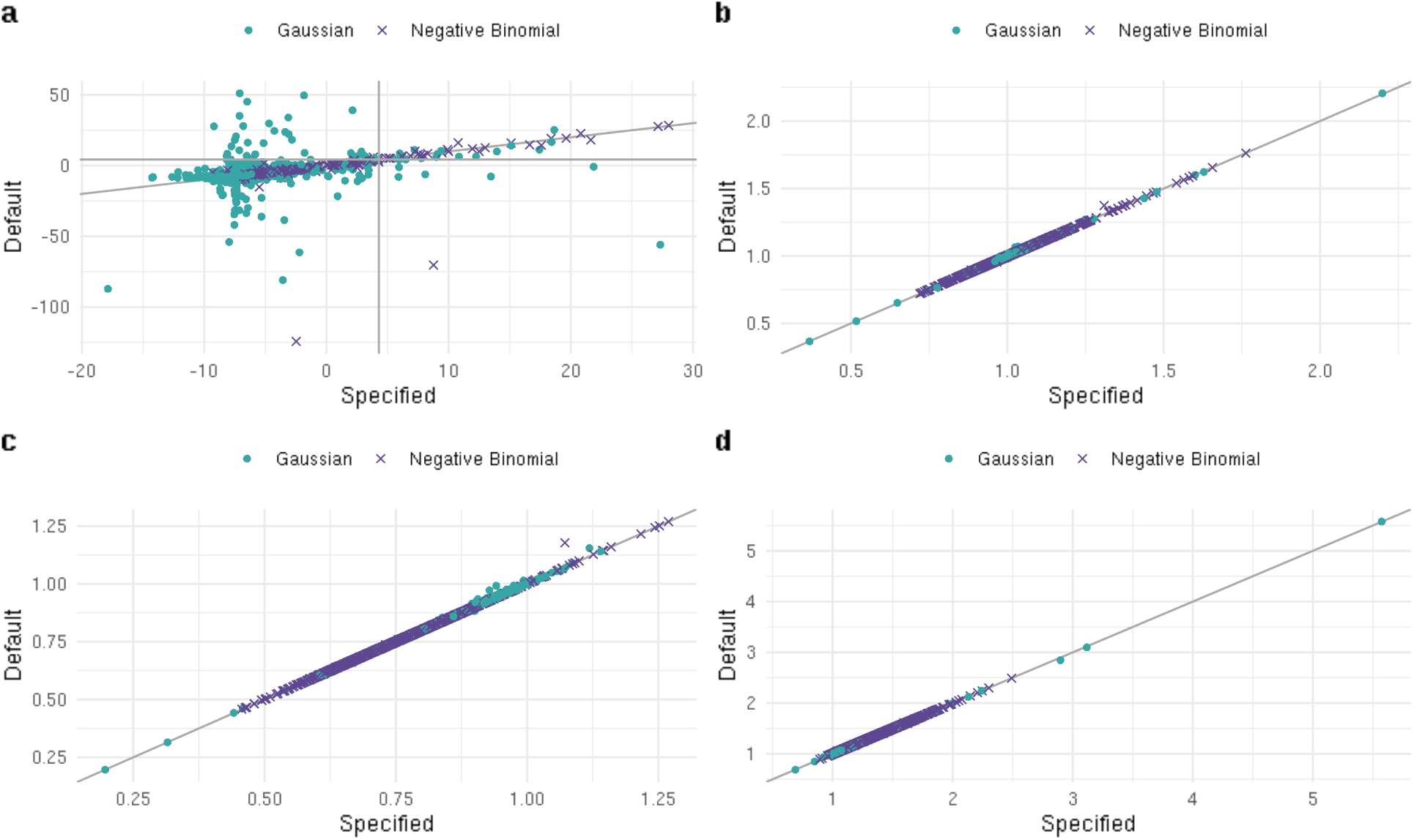
Comparing -log10 BF (figure a), fold change (figure b) and lower and upper credible interval bound (figure c and d, respectively) for BYM2 model under default prior or specified prior on INLA. We see that changing the prior can change the Bayes factor, specially negative values, but the fold change estimates remain the same.

**Supp Fig 7:**
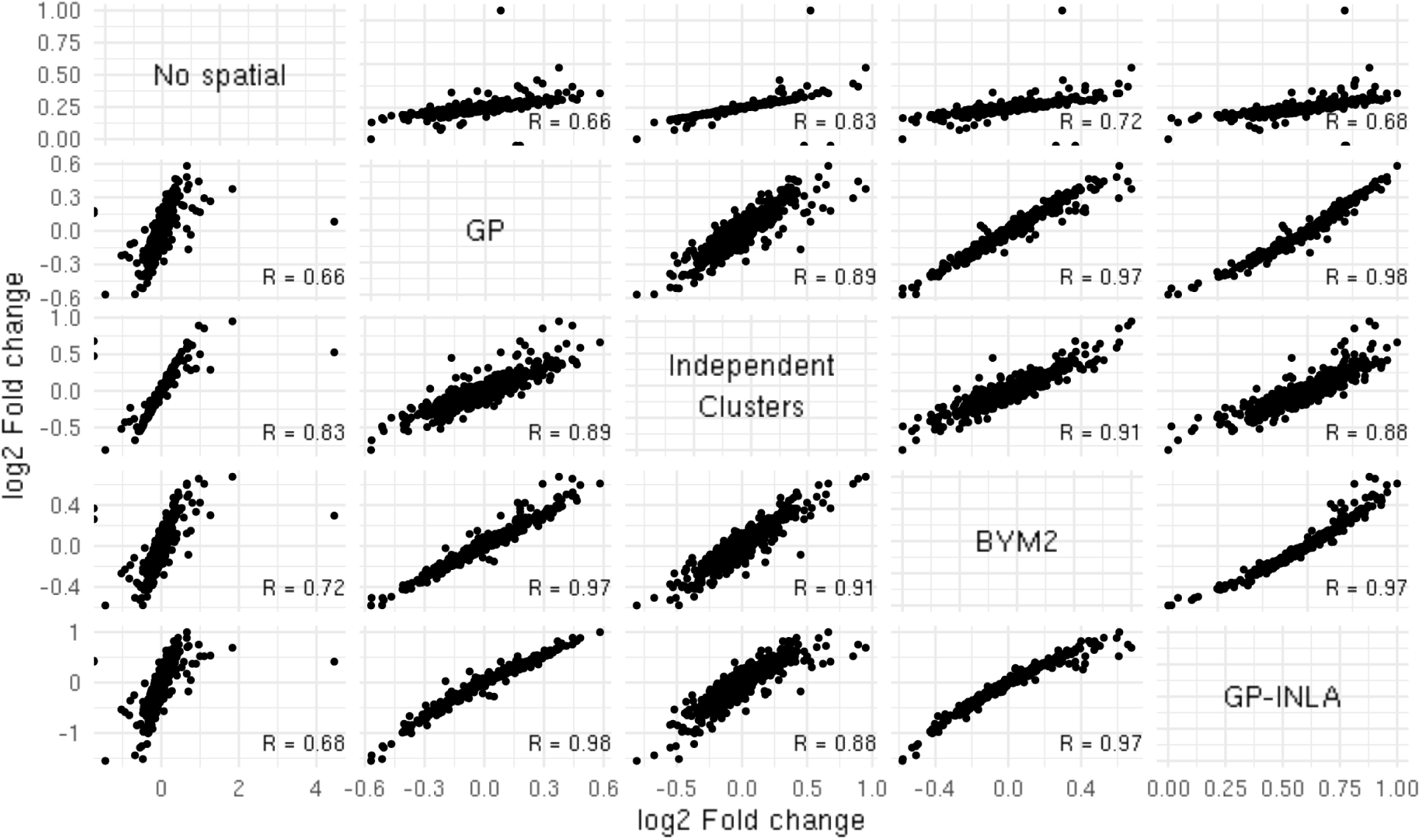
Comparing fold change estimates of different models assuming a Gaussian distribution while adjusting for transcript misassignment from segmentation error

**Supp Fig 8:**
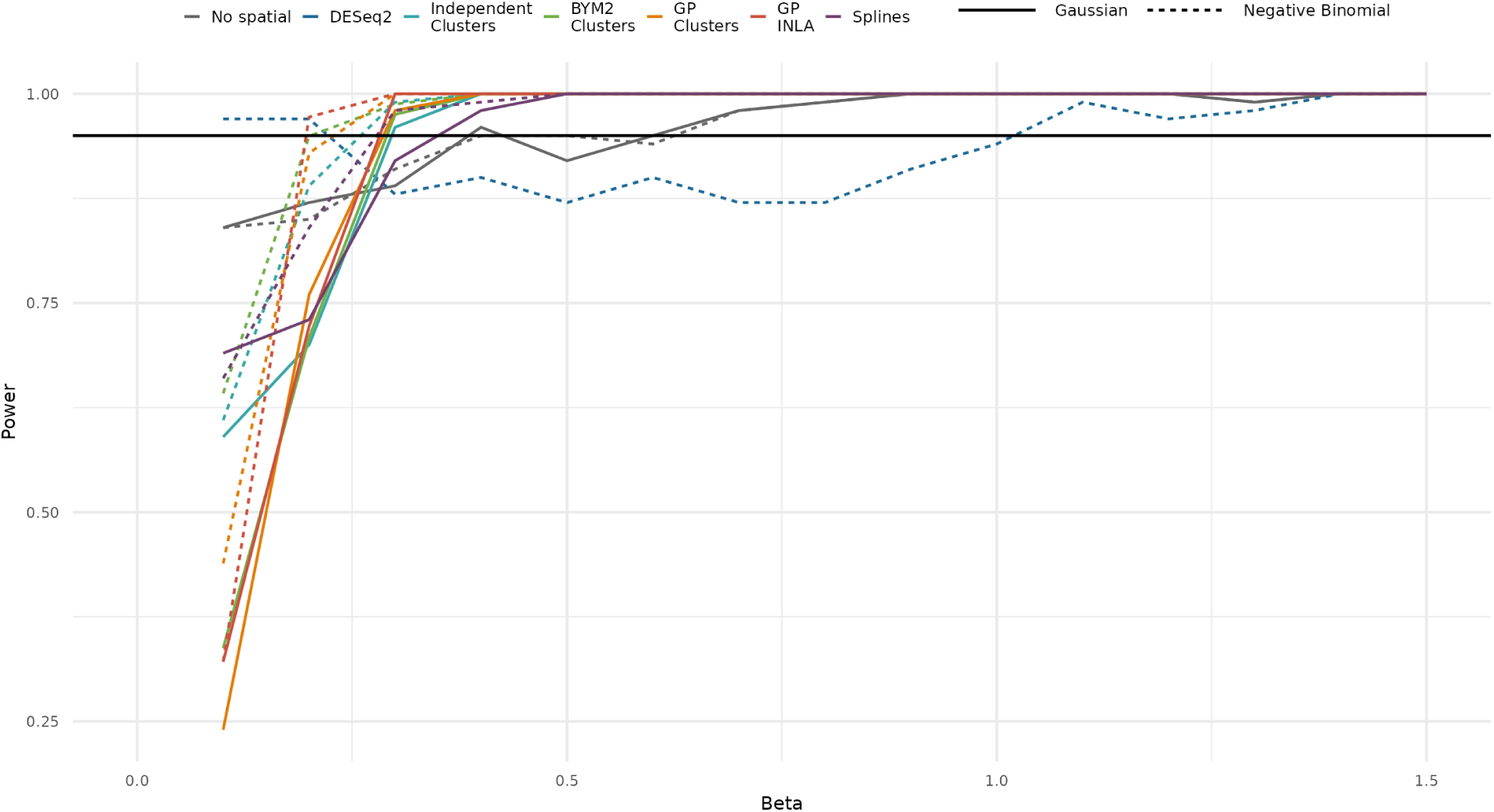
Power obtained based on results from repeated simulations. X axis indicates different niche coefficients, and y axis indicates power. Independent Clusters was considered with 5%n clusters and BYM2 and GP-INLA with 30%n. Due to inflates Type 1 error of models, Power is not directly comparable.

